# Novel genetic basis of resistance to Bt toxin Cry1Ac in *Helicoverpa zea*

**DOI:** 10.1101/2021.11.09.467966

**Authors:** Kyle M. Benowitz, Carson W. Allan, Benjamin A. Degain, Xianchun Li, Jeffrey A. Fabrick, Bruce E. Tabashnik, Yves Carrière, Luciano M. Matzkin

**Author notes:** Corresponding author: Kyle M. Benowitz. Department of Biology, Austin Peay State University, Sundquist Science Center, PO Box 4718, Clarksville, TN 37044.

## Abstract

Crops genetically engineered to produce insecticidal proteins from the bacterium *Bacillus thuringiensis* (Bt) have advanced pest management, but their benefits are diminished when pests evolve resistance. Elucidating the genetic basis of pest resistance to Bt toxins can improve resistance monitoring, resistance management, and design of new insecticides. Here, we investigated the genetic basis of resistance to Bt toxin Cry1Ac in the lepidopteran *Helicoverpa zea*, one of the most damaging crop pests in the United States. To facilitate this research, we built the first chromosome-level genome assembly for this species, which has 31 chromosomes containing 375 Mb and 15,482 predicted proteins. Using a genome-wide association study, fine-scale mapping, and RNA-seq, we identified a 250-kb quantitative trait locus (QTL) on chromosome 13 that was strongly associated with resistance in a strain of *H. zea* that had been selected for resistance in the field and lab. The mutation in this QTL contributed to but was not sufficient for resistance, which implies alleles in more than one gene contributed to resistance. This QTL contains no genes with a previously reported role in resistance or susceptibility to Bt toxins. However, in resistant insects, this QTL has a premature stop codon in a kinesin gene which is a primary candidate as a mutation contributing to resistance. We found no changes in gene sequence or expression consistently associated with resistance for 11 genes previously implicated in lepidopteran resistance to Cry1Ac. Thus, the results reveal a novel and polygenic basis of resistance.

## Introduction

Crops genetically engineered to produce insecticidal proteins from *Bacillus thuringiensis* (Bt) have provided control of some key pests during the past 25 years while reducing insecticide sprays and conserving arthropod natural enemies (Bravo *et al.* 2011; NASEM 2016; Dively *et al.* 2018; Romeis *et al.* 2018; Carrière *et al.* 2020a; Tabashnik *et al.* 2021). However, planting of a cumulative total of more than one billion hectares of Bt crops worldwide (ISAAA 2019) has selected for resistance that has reduced the efficacy of Bt crops against populations of at least nine major pest species (Calles-Torrez *et al.* 2019; Smith *et al.* 2019; Tabashnik and Carrière 2019). Knowledge of the genetic basis of pest resistance to Bt toxins can be useful for improving monitoring and management of resistance, as well as for designing new insecticides (Soberón *et al.* 2007; Jin *et al.* 2018).

Resistance to crystalline (Cry) Bt toxins typically entails mutations that reduce binding of the toxins to larval midgut receptors and thus block an essential step in the mode of action (Heckel *et al.* 2007; Peterson *et al.* 2017; Jurat-Fuentes *et al.* 2021). In particular, research has repeatedly implicated disruption or reduced expression of known or putative Bt toxin receptors from four protein families: ATP-binding cassette (ABC), cadherin, aminopeptidase N (APN), and alkaline phosphatase (ALP). Mutations that disrupt binding of toxins to receptors are frequently associated with high levels of resistance to one or a few closely related Bt toxins, weak or no cross-resistance to unrelated Bt toxins, and recessive inheritance of resistance (Mode 1 resistance; Tabashnik *et al.* 1998). Nonetheless, lepidopteran resistance to Bt toxins also includes examples where proteins from these families are not involved, toxin binding is not reduced, and inheritance of resistance is not recessive (Peterson *et al.* 2017; Jin *et al.* 2018).

Here we analyzed the genetic basis of resistance to Bt toxin Cry1Ac in the lepidopteran *Helicoverpa zea* (corn earworm or bollworm), which is one of the most economically important crop pests in the United States (Cook and Threet 2019; Musser *et al.* 2019). This polyphagous pest is the first insect reported to have evolved resistance to a Bt crop, specifically to cotton producing Cry1Ac (Luttrell *et al.* 1999; Ali *et al.* 2006; Tabashnik *et al.* 2008; Reisig *et al.* 2018). In contrast with Mode 1 resistance, inheritance of resistance to Cry1Ac in *H. zea* is not completely recessive (Brévault *et al.* 2013, 2015; Carrière *et al.* 2020b; Reisig *et al.* 2021). Caccia *et al.* (2012) concluded that resistance in their lab-selected AR1 strain of *H. zea* was complex, possibly polygenic, and not caused primarily by reduced binding of Cry1Ac to larval midgut membranes. Perera *et al.* (2021) found that knocking out the gene encoding the putative Cry1Ac receptor ABCC2 increased the concentration of Cry1Ac killing 50% of larvae (LC50) by 7- to 40-fold. Because >100-fold resistance to Cry1Ac is common in lab- and field-selected *H. zea* (Caccia *et al.* 2012; Brévault *et al.* 2013; Reisig *et al.* 2018; Kaur *et al.* 2019), they inferred that mutations disrupting ABCC2 are not the sole or primary mechanism of resistance to Cry1Ac in *H. zea.* Although most previous studies of Bt resistance in *H. zea* have emphasized the gene families listed above (Caccia *et al.* 2012; Zhang *et al.* 2019a; Fritz *et al.* 2020; Perera *et al.* 2021; Taylor *et al.* 2021), additional candidates have been identified using RNA-seq (Lawrie *et al.* 2020; 2022).

Our work focuses on the GA-R strain of *H. zea*, which had been selected for resistance to Bt toxins in the field and lab (Brévault *et al.* 2013, Welch *et al.* 2015). GA-R was derived from the moderately resistant GA strain, which had been selected for resistance to Bt toxins only in the field (Brévault *et al.* 2013). Relative to an unrelated susceptible lab strain (LAB-S) of *H. zea*, the LC50 of Cry1Ac was >500 times higher for GA-R and 55 times higher for GA (Brévault *et al.* 2013). Previous work identified reduced activation of Cry1Ac by midgut proteases as a potential field-selected mechanism of resistance that could explain part but not all of the resistance in GAR relative to LAB-S (Zhang *et al.* 2019a). Overall, the previous results with GA-R and other strains of *H. zea* summarized above led us to hypothesize that resistance to Cry1Ac in this species is polygenic and entails novel genetic mechanisms. Accordingly, genome-wide mapping approaches are warranted, but have been hindered because the only *H. zea* genome assembly available is highly fragmented (Pearce *et al.* 2017).

Here, we generated a chromosome-level genome for *H. zea*, then used a genome wide association study (GWAS), fine-scale mapping, and RNA-seq to elucidate the genetic basis of resistance to Cry1Ac in the GA-R strain of *H. zea.* We identified a quantitative trait locus (QTL) of 250 kb on chromosome 13 that was strongly associated with resistance to Cry1Ac. The results indicate a mutation in this QTL contributed to resistance but was not sufficient for resistance in GA-R. We also found no consistent association between resistance to Cry1Ac and any of 11 genes previously implicated in lepidopteran resistance to Bt toxins. We conclude the genetic basis of resistance to Cry1Ac in GA-R is novel and polygenic.

## Materials and methods

### Insect strains

We used four strains of *H. zea:* the highly resistant strain GA-R, its moderately resistant parent strain GA, the unrelated susceptible strain LAB-S, and the heterogeneous strain GA-RS that we created by crossing GA-R with LAB-S as described below. LAB-S was obtained from Benzon Research Inc. (Carlisle, PA, USA) and has not been exposed to Bt toxins or other insecticides. The resistant strain GA-R was derived from the third generation (F3) of the GA strain, which was started with 180 larvae collected on Cry1Ab corn from Tifton, Georgia in 2008 (Brévault *et al.*, 2013). GA-R was initially selected with Cry1Ac for nine generations and with MVPII thereafter (Brévault *et al.* 2013; Fritz *et al.* 2020; Carrière *et al.* 2020b). MVPII is a liquid formulation of a hybrid protoxin produced by transgenic *Pseudomonas fluorescens.* The amino acid sequence of the active portion of the protoxin is identical in the hybrid protoxin and Cry1Ac (Welch *et al.* 2015). For simplicity, hereafter we refer to MVPII as Cry1Ac. Amino acid sequence similarity between Cry1Ab and Cry1Ac is 86% (Carrière *et al.* 2015). Lab selection with Cry1Ac caused cross-resistance to Cry1Ab in GA-R (Welch *et al.* 2015) and in the AR strain of *H. zea* (Anilkumar *et al.* 2008). Moreover, adoption of Cry1Ac-producing cotton, a host plant of *H. zea*, was 94% in Georgia in 2008 (USDA 2008) and high in several preceding years (USDA 2020). Thus, the observed resistance to Cry1Ac in the GA strain (Brévault *et al.* 2013) could reflect direct selection in the field with Cry1Ac, cross-resistance from selection in the field with Cry1Ab, or both.

We conducted all rearing in walk-in growth chambers at 27 ± 1°C with 14h light: 10h dark photoperiod. We reared larvae on a casein-based wheat germ diet (Orpet *et al.* 2015a, 2015b) and conducted larval bioassays on Southland diet (Southland Products, Inc., Lake Village, AR, USA). We use these two different diets to optimize larval development and surface uniformity for toxin overlay in bioassays, respectively (Carrière *et al.* 2020b). Moths were kept in walk-in growth chambers under the same temperature and photoperiod mentioned above but under higher relative humidity than for larvae (60% Rh for moths and 20% Rh for larvae). Moths had access to a 10% sugar water solution for feeding and cheese cloth for oviposition (Welch *et al.* 2015). As previously reported (Fritz *et al.* 2020), we reared ca. 600 moths per generation for the first 35 and 33 generations of GA and GA-R, respectively. In 2012, we crossed GA with GA-R and used the resulting progeny to continue GA-R (Carrière *et al.* 2020b). After this interstrain cross, to reduce genetic drift and inbreeding, we reared two subsets of GA and crossed the two subsets every one to three generations (Carrière *et al.* 2020b). We used the same procedure to rear and cross two subsets of GA-R. Each subset had ca. 600 moths per generation (ca. 1200 moths per strain per generation). For GA, the mean number of moths per generation for F1 to F72 was ca. 900, based on the number of moths per generation of 600 for F1-F36 and 1200 for F37-72.

Relative to GA, GA-R had significantly higher survival on Bt cotton (producing Cry1Ac, Cry1Ac + Cry2Ab, or Cry1Ac + Cry1F) and Bt corn producing Cry1A.105 + Cry2Ab (Brévault *et al.* 2013, 2015; Carrière *et al.* 2018, 2019, 2020b, 2021). At the time we crossed GA-R with LAB-S in May 2018, we had selected GA-R with Cry1Ac for 58 generations. The GA-RS strain was created using mass reciprocal crosses between GA-R and LAB-S (i.e., 120 GA-R females × 120 LAB-S males and 120 LAB-S females × 120 GA-R males). GA-RS was founded with 450 neonates from each reciprocal cross. In the founding and subsequent generations, 900 larvae were reared and 600 adults (sex ratio 1:1) produced the neonates used for propagating the next generation.

### Genome sequencing of resistant strain GA-R

We generated a *de novo* genome assembly for GA-R using an approach combining a hybrid short- and long-read assembly with a long-read only assembly (Jaworski *et al.* 2020), which allows for improved error correction of long read data (Ye *et al.* 2016) without the need for massive coverage (Chakraborty *et al.* 2016). Hybrid assembly strategies have been used frequently with error-prone PacBio CLR data to generate highly contiguous genomes in non-model insect species (Hartke *et al*. 2019; Wan *et al.* 2019; Ferguson *et al*. 2020; Jaworski *et al.* 2020; Ma *et al.* 2020; Mathers 2020; Schmidt *et al.* 2020; Xu *et al.* 2021). For the short-read assembly, we collected and sequenced 30 GA-R larvae as described below. We trimmed reads and generated the assembly using Platanus (Kajitani *et al.* 2014). For the long-read assembly, we extracted DNA from the gut of a single GA-R fifth instar using a chloroform-based extraction method (Jaworski *et al.* 2020). PacBio libraries were constructed at the Arizona Genomics Institute (Tucson, AZ, USA). We then sequenced the library on a single lane of PacBio Sequel II, also at the Arizona Genomics Institute. We formatted raw PacBio reads using bam2fastq (https://github.com/jts/bam2fastq) and SeqKit (Shen *et al.* 2016) before filtering out all reads under 30-kb using Filtlong (https://github.com/rrwick/Filtlong). We mapped contigs from the short-read assembly to the long reads using DBG2OLC (Ye *et al.* 2016) before running three iterations of Sparc (Ye and Ma 2016) to correct the resulting contigs. We realigned the raw PacBio reads to the resulting assembly with pbmm2 (https://github.com/PacificBiosciences/pbmm2) and polished contigs using Arrow (Chin *et al.* 2013; https://github.com/PacificBiosciences/GenomicConsensus). Lastly, we mapped raw short reads to the assembly with Bowtie2 (Langmead and Salzberg 2012) to perform a final polishing step using Pilon (Walker *et al.* 2014).

We generated the PacBio-only assembly with Canu (Koren *et al.* 2017), using the reads longer than 30kb after filtering described above. We then polished the assembly using Arrow and Pilon as described above. Finally, we used Purge Haplotigs (Roach *et al.* 2018) to remove contigs containing alternate haplotypes generated due to high heterozygosity.

We aligned the two assemblies using nucmer within MUMmer4 (Marçais *et al.* 2018), keeping only alignments greater than 10 kb. We then generated the merged assembly using Quickmerge (Chakraborty *et al.* 2016). We performed additional merging by re-running Quickmerge with more liberal parameters on individual contig pairs after hypothesizing their contiguity based on synteny with *H. armigera* (Pearce *et al.* 2017; Valencia-Montoya *et al.* 2020). Specifically, chromosomes 5, 7, 8, 16, 17, 18, 19, 21, 23, 29, and 30 required such additional merging. After this step, only chromosome 17 had two contigs that did not merge. We therefore connected them with a default 100-bp gap according to NCBI standards (Karsch-Mizrachi *et al.* 2012). We again polished the final assembly using Arrow, Pilon, and Purge Haplotigs as above. Lastly, we ordered and named each chromosome to align with those of *H. armigera.* We generated a synteny plot comparing our genome to the *H. armigera* genome using Dot (https://github.com/marianattestad/dot) after filtering out alignments under 4000 bp in NUCmer.

We analyzed the completeness of the GA-R genome using BUSCO v.5 (Seppey *et al.* 2019), comparing genomic content against the lepidodptera_odb10 set of 5,286 conserved single copy genes. We calculated contig (before final merging of chromosome 17) and scaffold (final assembly) genome summary statistics with bbmap stats (https://sourceforge.net/projects/bbmap/). We calculated repeat content with RepeatModeler2 (Flynn *et al.* 2020) and RepeatMasker (Smit *et al.* 2013-2015). We provide a preliminary annotation produced following the funannotate pipeline (Palmer and Stajich 2020) with transcripts from *H. armigera* and proteins from *B. mori* used as evidence supporting putative annotations. We also used BUSCO v.5 to assess the completeness and accuracy of the annotation against the 5,286 single copy genes in the lepidoptera_odb10 dataset. To compare the quality of our assembly and annotation with the previous *H. zea* genome assembly (Pearce *et al.* 2017), we reanalyzed the genomic and proteomic BUSCO scores of that assembly against the same lepidoptera_odb10 dataset.

### Phenotyping of Cry1Ac-susceptible and -resistant larvae for genetic mapping

We used our standard diet overlay bioassay (Welch *et al.* 2015) to characterize GA-RS larvae as susceptible or resistant to Cry1Ac. We added 40 μl of a dilution containing 0.1% Triton X-100 and the desired concentration of Cry1Ac to the surface of solidified Southland diet in each well of 128-well bioassay trays (C-D International, Pitman, NJ, USA). One neonate (< 8 h old) was transferred to each well and trays were covered with ventilated plastic lids (C-D International) and held for 7 days under the abiotic conditions mentioned above.

We conducted five sets of bioassays using GA-RS neonates from the F2 (July 2018), F12 (July 2019), F22 (June 2020), F23 (July 2020), and F26 (October 2020) generations (Supplementary Table S1). Neonates were exposed to diet with 0 (control), 1, or 10 μg Cry1Ac per cm^2^ diet. After 7 days, live first instar larvae on diet with 1 μg Cry1Ac per cm^2^ were considered susceptible because this low toxin concentration inhibited their growth, whereas live larvae on diet with 10 μg Cry1Ac per cm^2^ that were third or subsequent instars were considered resistant because they grew well despite this high toxin concentration. For control larvae reared on diet without Cry1Ac, mean survival to third instar was 96% (range: 91-100%, mean n = 99 larvae per bioassay in five bioassays). Susceptible, resistant, and control larvae were transferred individually to plastic cups containing non-Bt diet, reared to fifth instar, transferred individually to 1.5 ml plastic tubes, and frozen at −80°C for subsequent genomic comparison.

### Genomic comparison of GA-R and LAB-S

We sequenced pools of larvae from the parental strains GA-R and LAB-S, which allowed us to evaluate genetic variation between and within these parental strains. This also allowed us to check if SNPs associated with resistance in the GWAS of GA-RS were more common in GA-R, and if those associated with susceptibility were more common in LAB-S. In April 2019, we collected 30 third instars from each strain, extracted DNA using Qiagen DNeasy Blood and Tissue Kits (Qiagen, Hilden, Germany), and constructed Illumina libraries using KAPA LTP Library Preparation Kits (Roche, Basel, Switzerland). We sequenced both libraries on an Illumina HiSeq4000 at Novogene (Beijing, China). We called variants with Platypus after read trimming and alignment to the genome using Trimmomatic and bwa-mem as described above. To detect potential selective sweeps in each strain, we used SAMtools mpileup (Li 2011) and PoPoolation (Kofler *et al.* 2011) to calculate Tajima’s D in 50-kb windows overlapping by 10 kb across the genome.

### Genome-wide association study for Cry1Ac resistance

From the heterogeneous strain GA-RS F12 larvae phenotyped in July 2019, we extracted DNA from 144 resistant and 144 susceptible larvae using ZYMO *Quick-DNA* 96 Plus Kits and quantified the DNA concentration of each individual using a Nanodrop (Thermo Fisher Scientific, Waltham, MA, USA). We combined equal amounts of DNA from each of the 144 resistant larvae to make a resistant pool and from each of the 144 susceptible larvae to make a susceptible pool, then generated libraries using KAPA LTP Library Preparation Kits for each pool. We sequenced both libraries on an Illumina HiSeq4000 at Novogene (Beijing, China).

We demultiplexed reads and trimmed for adapter contamination and low-quality sequence using Trimmomatic (Bolger *et al.* 2014). We mapped reads to the *de novo H. zea* genome using bwa-mem (Li and Durbin 2009). We used Platypus (Rimmer *et al.* 2014) to call and quantify SNP variants and short INDELs. We extracted biallelic SNPs with a minimum coverage of 20 in each pool and a total coverage between 60 and 500 from the Platypus output for statistical analysis, for which we used two approaches (Benowitz *et al.* 2019). First, we calculated a Z-statistic (Huang *et al.* 2012), using the formula 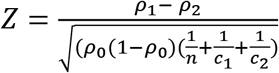, where and *ρ*_2_ are the reference alleles frequencies of each bulk, *ρ*_0_ is the mean allele frequency across bulks, *n* is the sample size of each bulk, and *c*_1_ and *c*_2_ are the read depth of each bulk. We evaluated statistical significance of *Z* against the standard normal distribution. Following convention (Barsh *et al.* 2012, Welter *et al.* 2014), we used *P* <5 × 10^-8^ as a threshold for significance. To estimate the density of significant sites, we used the R package WindowScanR (https://github.com/tavareshugo/WindowScanR) to calculate the percentage of SNPs with a more liberal threshold of *P* < 10^-5^ in 10-kb windows overlapping by 5 kb. Density of significant sites may be a particularly useful parameter because the magnitude of each individual *P*-value from a bulk segregant analysis is highly sensitive to noise. Close linkage to the causal allele, however, should result in a higher density of significant sites even if the *P*-value itself varies. Second, we performed a sliding-window analysis with 500-kb windows overlapping by 250 kb using the R package QTLseqr (Mansfeld and Grumet 2018), which implements the G′ method of Magwene *et al.* (2011). This method provides a statistical determination of whether an entire QTL, rather than an individual SNP, is significant, and also defines borders to QTLs deemed significant.

### Fine-scale mapping within chromosome 13

Using larvae from the F22 and F23 generation (July 2020), we conducted fine-scale mapping within the QTL in chromosome 13 associated with resistance, which we refer to hereafter as the *r1* locus. Using the methods described above, larvae were scored as resistant or susceptible.

After phenotyping, we reared susceptible larvae to fifth instars on untreated diet before storing all larvae (62 resistant and 51 susceptible) at −80°C. We extracted DNA from each sample using Qiagen DNeasy Blood and Tissue Kits and genotyped each larva individually via high-resolution melt curve (HRM) analysis at 12 SNP marker sites within the chromosome 13 QTL (Supplementary Table S2). For each site, we performed PCR in 10 μl reactions using Apex Taq DNA polymerase and EvaGreen Dye (Biotium, Fremont, CA, USA) as the intercalating dye. We ran RT-PCR in a QuantStudio 3 Real-Time PCR machine (Thermo Fisher) using continuous capture with a 0.025°C/s ramp. We used QuantStudio Design and Analysis Software (Thermo Fisher) to manually score melt curves for the identity of the focal SNP. We compared test melt curves against control curves generated from the parental LAB-S and GA-R strains. We used Fisher’s exact test to assess significant differences between allele frequencies of resistant and susceptible individuals.

The lack of amplification from some individuals for some markers yielded variation in sample size among the 12 markers. These ranged from 40 to 60 for resistant larvae (mean = 56) and 20 to 50 (mean = 45) for susceptible larvae. We also used HRM to obtain genotypes at marker #4 for 23 of 25 resistant larvae and 89 of 95 control larvae tested from the F26 generation. The control larvae were not screened with a bioassay and thus contained a mixture of resistant and susceptible individuals.

To confirm the accuracy of HRM genotyping, we Sanger sequenced a single site (marker 4; Supplementary Table S2) for all 60 resistant and a subset of 34 susceptible individuals. We designed new primers to produce a longer amplicon, and confirmed the quality of the resultant amplicons with gel electrophoresis. We cleaned the PCR product with Exonuclease I and Antarctic Phosphatase (New England BioLabs, Ipswich, MA, USA) before sending to Eurofins Genomics (Louisville, KY, USA) for Sanger sequencing.

In addition to the fine mapping data from the HRM, we used the data on significant SNP density from the original GWAS experiment as well as the Tajima’s D results from comparison of the GA-R and LAB-S strains to provide additional support for narrowing the region associated with resistance within chromosome 13.

### Inheritance and trajectory of resistance in GA-RS

We performed several analyses of genotype frequencies at marker 4 to understand how *r1* affects resistance. We used the genotypes from the RNA-seq study (see below) as a control group to examine the frequency of marker 4 in the GA-RS strain at the F26 generation. We also used a χ^2^ test to examine departure from Hardy-Weinberg equilibrium.

We used Fisher’s exact test to determine if resistant individuals from GA-RS were more likely to be homozygous for alleles from GA-R (GG) than heterozygous with one allele from GA-R and the other from LAB-S (GL) by comparing the frequencies of each genotype in the resistant samples from generations F22, F23, and F26 to the control samples from generation F26. We calculated the dominance parameter *h*, which varies from 0 for recessively inherited resistance to 1 for dominantly inherited resistance (Liu and Tabashnik 1997), using the genotype frequencies at marker 4 in the F22, F23, and F26 for resistant, control, and susceptible larvae (Supplementary Table S3). The results from F22 and F23 were similar and were pooled to increase the sample size for analyses.

To evaluate the relationship between genotype and resistance, we compared their trajectories across generations in GA-RS. We used linear regressions in R 4.1.0 to test the null hypothesis of no change in the log10 of percentage survival to third instar. Bioassays used 1 or 10 μg of Cry1Ac per cm^2^ in generations F2, F22 and F23, and only the higher concentration in the F26 test.

### RNA-seq and candidate gene analysis

To generate samples for RNA sequencing, we reared LAB-S, GA-R, and GA-RS (generation 26) individuals on untreated diet as described above in October 2020. When the larvae of the parental strains LAB-S and GA-R reached the third instar, we dissected midguts from 15 individuals and froze them in groups of five, generating three biological replicates per strain. For the GA-RS heterogeneous strain, we dissected 95 third instar midguts and froze them individually, while simultaneously freezing the remainder of the carcass separately in wells of a PCR plate. We extracted DNA from each sample using a squish extraction (Gloor *et al.* 1993) using 50 μl of buffer. We screened samples HRM as above at marker 4, which was one of two sites we found to be most strongly associated with resistance. We selected 15 individuals that were homozygous for the LAB-S genotype at this site (henceforth “LL”) and 15 individuals homozygous for the GA-R genotype at this site (henceforth “GG”) and pooled the midguts corresponding to these samples into groups of five, again generating three biological replicates for each genotype. Selecting genotypes in this way allowed us to isolate the effects that the chromosome 13 QTL has on gene expression, giving us the potential to detect *trans*-regulatory effects. We extracted RNA from all 12 midgut samples using ZYMO Direct-zol RNA Miniprep Kit kits and built paired-end libraries with KAPA stranded mRNA seq kits. Libraries were sequenced in part on an Illumina NextSeq at the University of Arizona Genetics Core (UAGC; Tucson, AZ, USA) and part on an Illumina NovaSeq at Novogene.

We trimmed reads using Trimmomatic (Bolger *et al.* 2014) and aligned them to our *H. zea* assembly with Hisat2 (Kim *et al.* 2019). We built genome-guided transcriptome assemblies for each sample using StringTie (Pertea *et al.* 2016) and used StringTie merge to create a single transcriptome. We used blastp to identify the closest ortholog in *H. armigera* for each gene. We quantified read abundance for each sample using Salmon (Patro *et al.* 2017) and combined its transcript-level counts into gene-level counts with the R package tximport (Soneson *et al.* 2016). We analyzed differential expression using FDR-corrected *P*-values from negative binomial models at the gene level with edgeR (Robinson *et al.* 2010), after filtering and normalization for library-size differences. We performed statistical comparisons of LAB-S to GA-R and LL to GG. We performed a one-tailed (to account for directionality of gene expression differences) χ^2^ test to examine whether the overlap of differentially expressed (DE) genes was more than expected by chance.

In addition to analyzing global differential expression, we used the RNA-seq data to better understand gene structure and expression within the candidate QTL. For each sample, we used Trinity (Haas *et al.* 2013) to construct *de novo* transcriptome assemblies for each sample. Using tblastn, we identified the transcripts in each Trinity assembly corresponding to all StringTie transcripts from the region from bp 4,370,000 – 4,620,000. We then took the longest transcript from each Trinity assembly and used orfipy (Singh and Wurtele 2021) to extract the longest open reading frame (ORF) for each gene. Next we compared the ORFs from each of the 12 samples, looking for differences in predicted protein structure between the samples with resistant chromosome 13 genotypes (GA-R and GG) and susceptible ones (LAB-S and LL). We quantified midgut abundance of transcripts for all genes expressed in this region with average log count per million reads across all samples produced by edgeR. We used PROVEAN (Choi and Chan 2015) to predict the effects of amino acid substitutions between LAB-S and GA-R for each of the genes in this region.

After identifying a nonsense mutation in the *kinesin-12* gene, we manually inspected this site in IGV (Robinson *et al.* 2017). After performing Sanger sequencing of the GA strain and field samples (see below), we additionally inspected this site by visualizing sequencing chromatograms in Teal (Rausch *et al.* 2020). We analyzed the putative kinesin-12 protein computationally using blastp to find homologous proteins. We then used blastp to compare the sequence conservation upstream and downstream of the stop codon mutation in the lepidopteran species *H. armigera* (XP_021193241.1), *Chloridea virescens* (PCG76683.1), *Spodoptera litura* (XP_022828947.1), *Manduca sexta* (KAG6448083.1), and *Bombyx mori* (XP_004927959). We used a two-tailed paired t-test to assess amino acid conservation across these species before and after the nonsense mutation. We aligned these sequences with both the GA-R and LAB-S *H. zea* sequences with Clustal Omega (Sievers and Higgins 2018) and plotted the alignments using TeXshade (Beitz 2000). To further probe the potential structure and function of the protein, we used InterProScan 5 (Jones *et al.* 2014) to search for additional protein domains, AlphaFold 2.1.0 (Jumper *et al.* 2021) to predict the 3D protein structure, SignalP 5.0 (Armenteros *et al.* 2019) to examine if the protein contained signal peptides, and DeepGOPlus (Kulmanov and Hoehndorf 2020) to predict gene ontology (GO; Ashburner *et al.* 2000) categorization based on the sequence.

### Genotyping of the GA strain and field samples for the *kinesin-12* mutation

We collected *H. zea* larvae from the field in Maricopa, Arizona in October 2020 and Tifton, Georgia in July 2021. Both populations had high resistance to Cry1Ac (Yu *et al.* 2021; Y. Carrière, unpubl. data). We reared larvae to adults in the lab and collected tissue from 25 of the Maricopa samples and 39 of the Tifton samples. We extracted DNA using a squish extraction (Gloor *et al.* 1993) in 150 μl of buffer. We analyzed five F72 GA individuals sequenced in October 2016 (Fritz *et al.* 2020). We downloaded raw reads from NCBI (PRJNA599999), trimmed them using Trimmomatic, aligned them to the GA-R genome with bwa-mem, and identified the frequency of the C546T mutation with samtools and VarScan. We additionally collected tissue from 20 F87 adults from the GA strain in May 2018 and extracted DNA with Qiagen DNeasy Blood and Tissue Kits. We designed primers (Supplementary Table S2) to amplify the region surrounding the *kinesin-12* mutation causing a stop codon. We performed PCR as above, although with the addition of final concentration 0.1 μg/μl bovine serum albumin (Sigma-Aldrich, St. Louis, MO, USA) due to PCR inhibitors. Sanger sequencing was done at Eurofins Genomics as described above. We screened sequences manually in Geneious Prime (Biomatters, Auckland, NZ) for presence of the target mutation.

### Analysis of 11 genes previously implicated in lepidopteran resistance to Cry1Ac

We used our results from GWAS and RNA-seq to test the hypothesis that one or more of 11 genes previously implicated in lepidopteran resistance to Cry1Ac contributed to resistance in our strains. For each gene, we present the lowest *P*-value from the original GWAS of SNPs between the start and end of the gene in the G′ analysis. We also report the FDR corrected *P*-values from differential expression analyses in edgeR comparing GA-R versus LAB-S and within GA-RS comparing individuals with both alleles from GA-R (GG) versus those with both alleles from LAB-S (LL).

## Results

### Chromosome-level assembly of the genome of resistant strain GA-R of *H. zea*

We generated a *de novo* chromosome-level assembly of the genome of the GA-R strain of *H. zea* with 31 chromosomes, 375.2 Mb, 36.9% GC content, 33.0% repeat content, an N50 scaffold length of 12.9 Mb, and 15,482 encoded proteins (Supplementary Table S4). Of 5,286 conserved single-copy lepidopteran genes, this genome has 98.9% complete, 98.5% complete and single-copy, 0.4% duplicated, 0.3% fragmented, and only 0.8% missing. The new genome assembly has only one gap set to 100 bp, which occurs in chromosome 17 and represents 0.000027% of the genome. This is a considerable improvement from the *H. zea* assembly of Pearce *et al.* (2017), which has 34.1 Mb of gaps representing 10% of that genome (Supplementary Table S4). Relative to the previous *H. zea* genome assembly, the new assembly has 64-fold greater N50, 10% more base pairs, 15% more complete BUSCO genes, and double the repeat content (Supplementary Table S4). Relative to previous estimates based on bacterial artificial chromosome sequencing and flow cytometry, the new genome size is 3% larger than an estimate for *H. zea* (Coates *et al.* 2017) and 5% smaller than an estimate for *H. armigera* (Zhang *et al.* 2019b).

The 31 chromosomes in the new assembly are largely syntenic with those of *H. armigera* (Pearce *et al.* 2017; Valencia-Montoya *et al.* 2020), although with different inversion karyotypes for 19 chromosomes (Supplementary Figure S1). We also found substantial differences between *H. zea* and *H. armigera* in the Z chromosome (chromosome 1) that are not associated with reversed sequences and thus probably not caused by chromosomal inversions (Supplementary Figure S1). An alternative hypothesis is that errors in one or both assemblies contributed to these differences. Errors are less likely in the new *H. zea* assembly because of its higher N50 and lower gap percentage relative to the *H. armigera* assembly (Supplementary Table S4, Pearce *et al.* 2017). In particular, the Z chromosome assembled cleanly without additional merging in the new *H. zea* assembly.

### Genomic comparison between GA-R and the susceptible strain LAB-S

Sequencing of 30 larvae from GA-R and 30 larvae from the unrelated susceptible strain LAB-S revealed 165,416 fixed differences between the strains, as well 941,146 variable sites in GA-R and 911,946 in LAB-S. Analysis of Tajima’s D showed many regions with low genetic variation throughout the genome in both strains (Supplementary Figures S2 and S3). For both strains, these regions could reflect genetic drift or selective sweeps caused by adaptation to lab conditions. For GA-R, regions of low variation could also reflect selection for resistance in the lab.

### Genome-Wide Association Study (GWAS) of resistance to Cry1Ac in GA-RS

We created the GA-RS strain by crossing GA-R and LAB-S. Both the Z-score and G′ sliding window analyses of 1,578,733 SNPs from the GWAS using pools of resistant and susceptible larvae from the F12 generation of GA-RS identified a region associated with resistance on chromosome 13 (Figure 1, Supplementary Figure S4). G′ analysis via QTLseqr defined this QTL as a region from 4.0 to 6.5 Mb. This QTL contains 117 SNPs associated with resistance at *P* < 5 × 10^-8^. All of these 117 SNPs showed the expected relationship with the parental strain. The 108 alleles found at higher frequency in the resistant larvae were more common in GA-R than LAB-S. The remaining nine alleles were at higher frequency in the susceptible larvae and were more common in LAB-S than GA-R.

**Figure 1.**
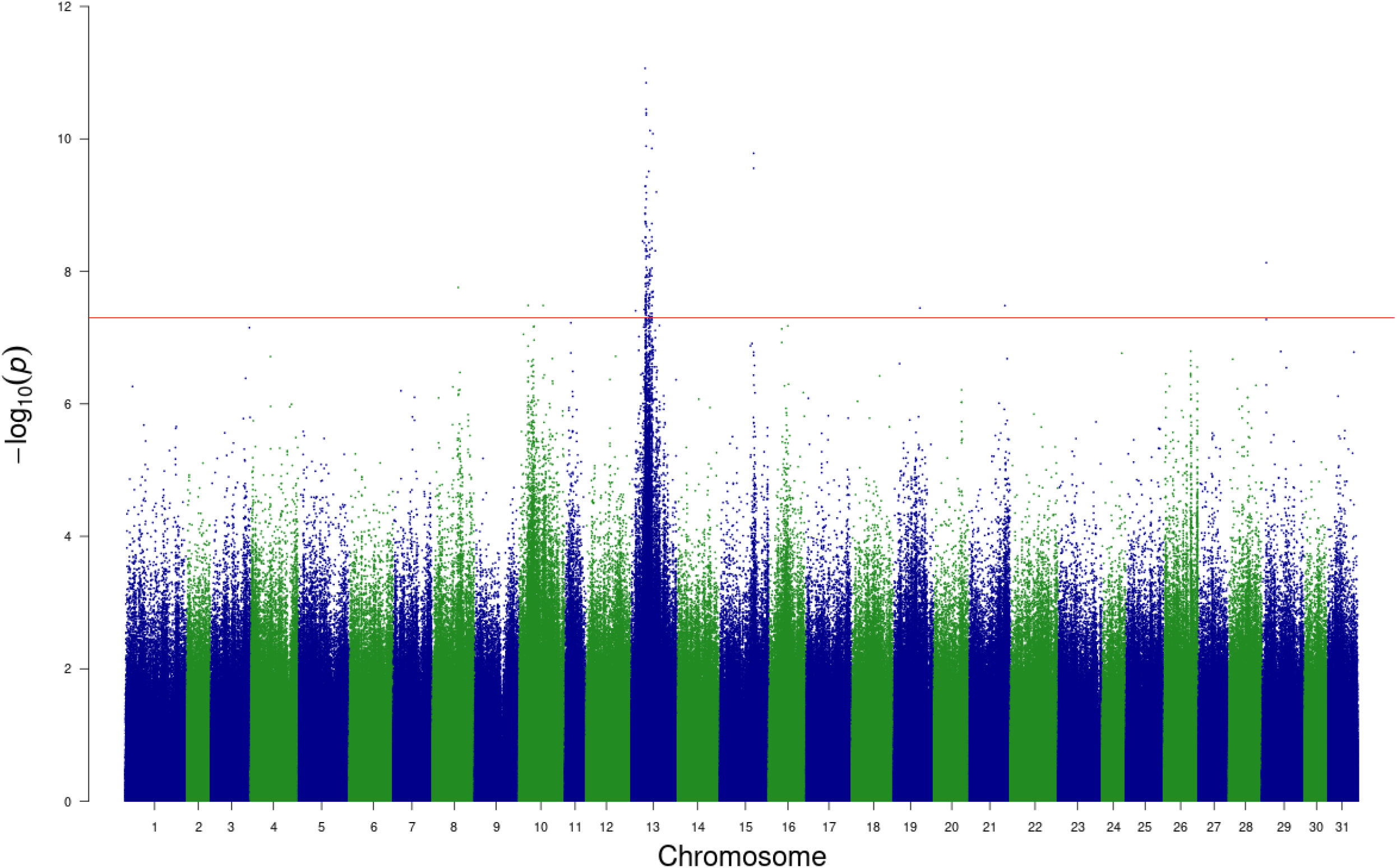
Manhattan plot of GWAS results showing –log_10_ of the *P*-values for Z-scores comparing allele frequencies between resistant and susceptible larvae. The red line indicates the threshold for significant association (*P* = 5e^-8^).

Outside of this QTL, only 18 SNPs were associated with resistance at *P* < 5 ×10^-8^. Both chromosome 10 and 15 had two of these SNPs. No other chromosome had more than one. In the G′ analysis, only the QTL on chromosome 13 was significantly associated with resistance (Supplementary Figure S4). We refer to this QTL as the *r1* locus.

Analysis of Z-scores from the GWAS shows that SNP sites that differed significantly between resistant and susceptible larvae were not evenly distributed across the chromosome 13 QTL (Figure 2A). Eighteen of the 25 windows of 10 kb with the top 5% significant SNP density were clustered between 4.42 and 4.60 Mb. Consistent with the GWAS results, Tajima’s D provides evidence for a selective sweep in GA-R between 4.3 and 5.2 Mb on chromosome 13 (Figure 2B).

**Figure 2.**
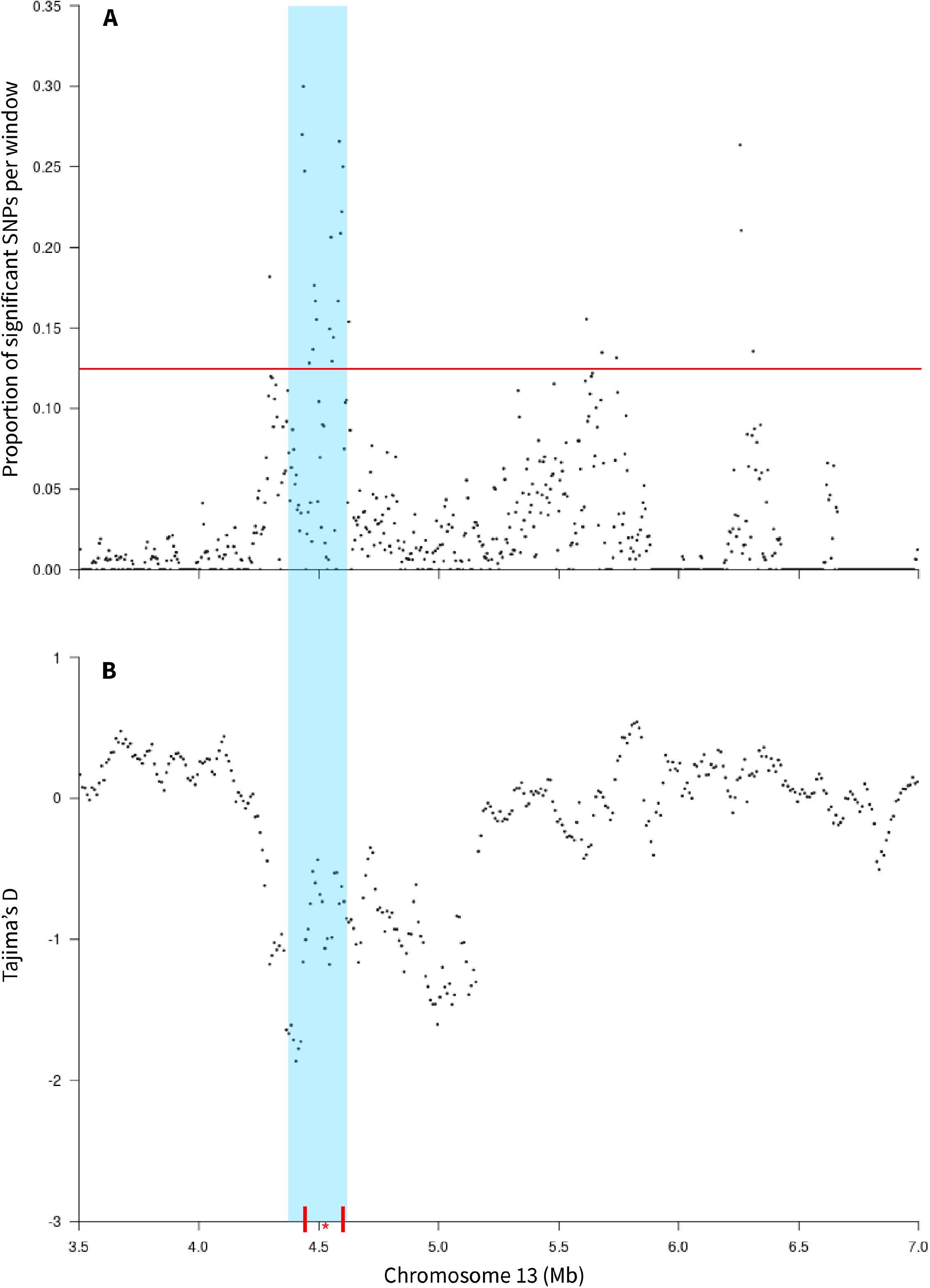
Association between resistance to Cry1Ac and SNPs within the chromosome 13 QTL. (A) Proportion of significant SNPs (*P* < 1e^-5^) from the Z-score analysis of the QTL data in 10-kb sliding windows. The horizontal red line indicates the 95^th^ percentile of the distribution. (B) Evidence of a selective sweep in GA-R from Tajima’s D in 50-kb sliding windows. Blue shading covers the QTL from 4.37 to 4.62 Mb. The vertical red bars show the locations of markers 4 and 5 (Table 1). The red asterisk indicates the location of *kinesin-12.*

### Fine-scale mapping within the *r1* locus

We used HRM to genotype individual resistant and susceptible larvae from the F22 and F23 of GA-RS for SNPs at 12 marker sites within *r1.* This revealed eight markers (2 to 9) from 4.3 to 5.0 Mb significantly associated with resistance (Table 1). For marker 4 at 4.5 Mb (n = 57) and marker 5 at 4.6 Mb (n = 60), all resistant larvae genotyped from GA-RS were either homozygous for the allele from the resistant GA-R strain (GG) or heterozygous, with one allele from GA-R and the other from the susceptible LAB-S strain (GL) (Table 1). These results were confirmed via Sanger sequencing for all 60 resistant individuals and 34 susceptible individuals. A similar test using only resistant larvae from the F26 confirmed this result: all 23 resistant larvae genotyped were either GG (16) or GL (7). By contrast, the three genotypes were in Hardy-Weinberg equilibrium in 89 larvae genotyped from a control sample from the F26 that was not screened for resistance and thus contained a mixture of resistant and susceptible individuals (24 GG: 44 GL: 21 LL, χ^2^ = 0.0091; *P* = 0.52). Genotype frequency in the F26 larvae differed significantly between the resistant larvae and the control larvae (χ^2^ = 22.60, *P* = 1.2 × 10^-5^).

**Table 1.** Genotype and allele frequencies at 12 markers on the chromosome 13 QTL for resistant and susceptible GA-RS larvae from generations F22 and F23.

| Marker | Bp (Chr13) | Resistant larvae |  | Susceptible larvae |  | *P-value*<sup>b</sup> |
| --- | --- | --- | --- | --- | --- | --- |
|  |  | Genotypes<br>(GG / GL / LL)<sup>a</sup> | G allele<br>frequency | Genotypes<br>(GG / GL / LL)<sup>a</sup> | G allele<br>frequency |  |
| 1 | 4,110,560 | 41 / 15 / 4 | 0.81 | 23 / 23 / 3 | 0.70 | 1.0 |
| 2 | 4,281,434 | 30 / 23 / 7 | 0.69 | 9 / 24 / 16 | 0.43 | 9.6 e-03 |
| 3 | 4,379,538 | 41 / 18 / 1 | 0.83 | 13 / 24 / 12 | 0.51 | 2.4 e-04 |
| 4 | 4,475,706 | 40 / 17 / 0 | 0.85 | 13 / 24 / 12 | 0.51 | 4.0 e-05 |
| 5 | 4,596,970 | 43 / 17 / 0 | 0.86 | 13 / 24 / 12 | 0.51 | 3.0 e-05 |
| 6 | 4,720,567 | 39 / 17 / 2 | 0.82 | 12 / 13 / 12 | 0.50 | 1.3 e-04 |
| 7 | 4,829,519 | 40 / 18 / 1 | 0.83 | 13 / 23 / 13 | 0.50 | 1.0 e-04 |
| 8 | 4,902,478 | 38 / 21 / 1 | 0.81 | 13 / 23 / 13 | 0.50 | 1.0 e-04 |
| 9 | 4,998,799 | 38 / 19 / 1 | 0.82 | 12 / 23 / 14 | 0.48 | 5.0 e-05 |
| 10 | 5,226,319 | 27 / 19 / 3 | 0.75 | 14 / 24 / 5 | 0.61 | 0.47 |
| 11 | 5,727,295 | 11 / 22 / 7 | 0.55 | 5 / 14 / 1 | 0.60 | 0.29 |
| 12 | 6,232,397 | 15 / 19 / 8 | 0.58 | 3 / 33 / 8 | 0.44 | 1.0 |

<sup>a</sup> G indicates the allele was more common in the resistant GA-R strain,
L indicates the allele was more common in the susceptible LAB-S strain.
<sup>b</sup> From Fisher's exact test of the null hypothesis that allele frequency did not differ between the resistant and susceptible larvae.

The results from the GWAS, Tajima’s D, the G′ analysis, analysis of SNP density, and fine-scale mapping (Figures 1 and 2, Table 2), identify the region between 4.3 to 4.6 Mb as most likely to contain the mutation(s) causing the effect of chromosome 13 on resistance to Cry1Ac. This region is captured by a single contig in both our Canu and DBG2OLC assemblies (Supplementary Figure S5) and is syntenic with a region of the *H. armigera* chromosome 13 (Supplementary Figures S1 and S6).

**Table 2.** Larval midgut expression of genes in the region of chromosome 13 QTL associated with resistance to Cry1Ac

| Gene ID | Start - Stop (orientation) | Name | Larval midgut expression (mean log<sub>2</sub> CPM) |
| --- | --- | --- | --- |
| MSTRG.8092 | 4,371,598 – 4,491,148 (+) | *Cyclic AMP-response element-binding protein A* | 2.44 |
| MSTRG.8095 | 4,491,241 – 4,520,601 (-) | *Heparan-alpha-glucosaminide-N-acetyltransferase* | -0.02* |
| MSTRG.8097 | 4,504,492 – 4,507,681 (+) | *Juvenile hormone esterase-like Carboxyl/cholinesterase CCE006D* | 4.84 |
| hz_G0000107 | 4,536,278 – 4,541,251 (+) | *Lipase member H-B-like* | N/A |
| MSTRG.8100 | 4,541,250 – 4,593,328 (-) | *Phosphatidylinositol 4-phosphate 3-kinase C2 domain-containing subunit alpha* | 3.25 |
| MSTRG.8101 | 4,546,116 – 4,547,856 (+) | *Kinesin-related protein 12-like* | 7.08 |
| hz_G0000111 | 4,593,482 – 4,595,264 (-) | Uncharacterized protein | N/A |
| hz_G0000112 | 4,600,520 – 4,606,939 (+) | Uncharacterized protein | N/A |
| MSTRG.8102 | 4,607,861 – 4,617,975 (-) | *Ubiquitin-protein ligase E3A* | 3.88 |
| MSTRG.8103 | 4,618,419 – 4,620,800 (+) | *Retinal rod rhodopsin-sensitive cGMP 3', 5'-cyclic phosphodiesterase subunit delta* | 2.98 |

Gene IDs refer to the StringTie annotation when expressed for correspondence with the RNA-seq data. Funannotate IDs refer to non-expressed genes. N/A indicates the gene was not expressed. \*Because of the low expression of this gene (12-fold lower than the median for the other six expressed genes), we considered it not to be substantially expressed.

### Gene expression in the midgut and a stop codon in *r1*

Based on the results above and annotations from funannotate and StringTie, we focused on the 10 genes between 4.37 and 4.62 Mb on chromosome 13. Six of these 10 genes were expressed substantially in the midgut of third instar larvae (Figure 3A, Table 2). The most highly expressed gene encodes a wild-type protein of 308 amino acids that has sequence identity of 97% with *kinesin-related protein 12* in *H. armigera* (XP_021193241; Supplementary Figure S7, Supplementary Table S5). The structure of this gene in terms of introns and exons is also similar between *H. zea* and *H. armigera* (Supplementary Figure S6B). Hereafter, we refer to this gene in *H. zea* as *kinesin-12*.

**Figure 3.**
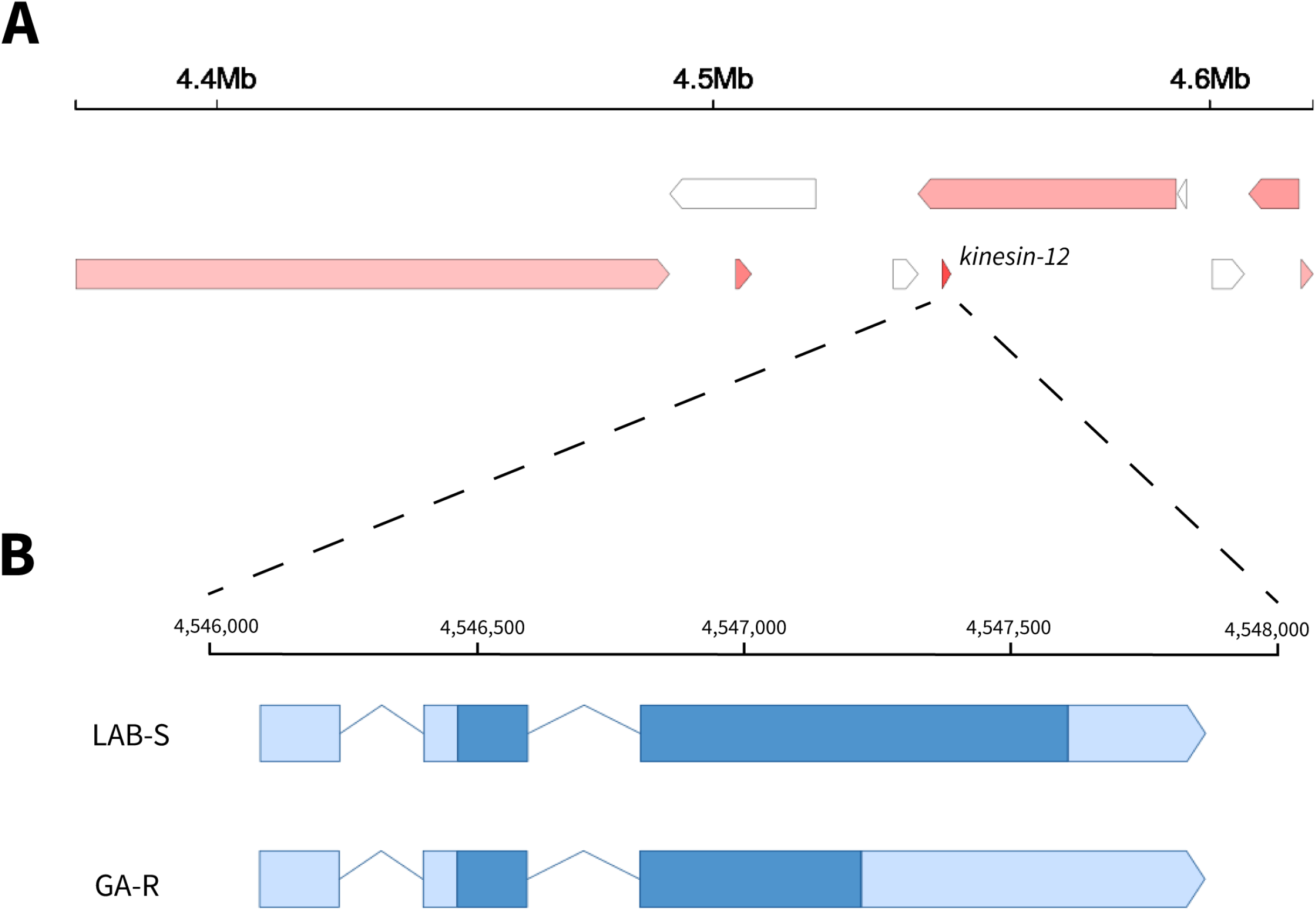
Ten genes including *kinesin-12* in the resistance-associated QTL on chromosome 13. (A) The four genes at the top are in the (–) orientation, the other six below are in the (+) orientation (Table 2). The four genes in white were not expressed substantially in the midgut. Darker red indicates higher expression in the midgut (Table 2). (B) The structure of *kinesin-12* in LAB-S and GA-R. Boxes represent exons, light blue indicates UTRs, and dark blue signifies coding regions.

In GA-R we found a point mutation (C546T) in *kinesin-12* that introduces a premature stop codon expected to truncate the protein at 183 amino acids (Figure 3B, Supplementary Figure S7). Manual inspection of aligned genomic reads, RNA-seq reads, and Sanger sequences further confirmed the identity of the SNP (Supplementary Figures S7, S8 and S9). This mutation occurred in 100% of reads covering the SNP from 30 GA-R larvae and in 0% of reads covering the SNP from 30 LAB-S larvae that were sequenced in the genomic comparison between these strains. In the GWAS with GA-RS, this mutation was more common in resistant larvae (71%) than susceptible larvae (32%; *P* = 7.48 × 10^-6^). It occurs at bp 4,547,246, between the two markers (4 and 5) most tightly associated with resistance to Cry1Ac (Table 1). Furthermore, RNA-seq near marker 4 detected the C546T mutation in 100% of reads covering the SNP from 15 GG larvae and 0% of reads covering the SNP from 15 LL larvae, confirming complete linkage between this mutation and marker 4. All of this evidence identifies the C546T mutation in *kinesin-12* as a candidate for causing the contribution of the *r1* allele to resistance to Cry1Ac.

Aside from the *kinesin-12* mutation, we detected missense mutations between GA-R and LAB-S linked to marker 4 in three of the other six candidate genes in this region that were expressed substantially in midguts of third instar larvae. These genes encode juvenile hormone esterase, phosphatidylinositol 4-phosphate 3-kinase, and ubiquitin protein ligase (Table 2). However, according to PROVEAN, none of the amino acid substitutions in these genes are expected to have major effects on protein function.

Although the *kinesin-12* gene has been annotated as encoding a kinesin-related protein in *H. armigera*, both its wild-type function and the effects of the nonsense mutation remain unclear. Within the moth family Noctuidae, amino acid sequence identities relative to the LAB-S strain of *H. zea* are 97% for *H. armigera* (as noted above), 87% for *C. virescens*, and 61% for *S. litura* (Supplementary Table S5). Outside this family, no annotated proteins in Lepidoptera have greater than 45% amino acid identity and we found no orthologs in other insect orders. For five species of Lepidoptera, including the three mentioned above plus *B. mori* and *M. sexta*, the sequence identity for this protein relative to LAB-S did not differ significantly between upstream and downstream from the stop codon (t_4_ = 0.95, *P* = 0.40; Supplementary Table S5). Thus, we cannot reject the null hypothesis that evolutionary constraints are similar for the portions of the protein before and after the stop codon.

Analysis with SignalP found no evidence for a signal peptide, indicating this protein is not likely to be integrated into or secreted through the cell membrane. The most specific of the GO terms associated with this protein by DeepGoPlus (Supplementary Table S6) is intracellular non-membrane-bounded organelle, which is most closely associated with kinesin-related proteins in *Drosophila melanogaster* (http://amigo.geneontology.org/amigo/term/GO:0043232). InterProScan and AlphaFold identified a coil with high confidence (aa 124-194; Supplementary Figures S11 and S12) that would be disrupted in the truncated form of the protein. Thus, we find moderate evidence the *H. zea kinesin-12* gene encodes a kinesin protein whose function might be disrupted by the C546T nonsense mutation.

### *Kinesin-12* mutation in the GA strain and in field samples

To test the hypothesis that the C546T mutation in *kinesin-12* originated in the field, we determined its frequency in the GA strain of *H. zea*, which was selected for resistance in the field, but not in the lab (Brévault *et al.* 2013; Welch *et al.* 2015). In GA, the frequency of the C546T mutation was 0.80 in five larvae from the F72 generation (three with homozygous TT genotypes and two with heterozygous CT genotypes), which does not differ significantly from its frequency of 0.975 in 20 larvae from the F87 generation (19 homozygous TT and one heterozygous CT; Fisher’s exact test: *P* = 0.10). These results are consistent with the hypothesis that C546T mutation originated in the field population from which GA was established. The alternative hypothesis that this mutation was absent in the field and arose in the lab seems unlikely. Based on the mean of ca. 900 moths per generation for GA and assuming a mutation rate of 3 × 10^-9^ per nucleotide site (Keightley *et al.* 2015; Yoder and Tiley 2021), the probability of a single mutation arising at a particular nucleotide site in GA during 72 generations is 0.0004.

The high frequency of C546T in GA after rearing for many generations in the lab without exposure to Bt toxins implies this mutation caused little or no fitness cost when reared in the lab in the absence of Bt toxins. However, we did not find this mutation in 39 individuals collected from the field in Georgia in July 2021 or in 25 individuals derived from the field in Arizona in 2020, despite the resistance to Cry1Ac in both of these field populations (Yu *et al.* 2021; Y. Carrière, unpubl. data). Whereas all individuals from the Georgia 2021 sample had the same sequence as LAB-S at the codon starting with bp 546, three individuals from Arizona had a single base pair substitution (C546A) changing the encoded amino acid from glutamine to lysine. In the field samples from Arizona in 2020 and Georgia in 2021, we detected no insertions, deletions, or other mutations introducing a stop codon in the 200 bp upstream or downstream from the C546T mutation.

### Inheritance and trajectory of resistance in GA-RS

The genotype frequencies at marker 4 in resistant and susceptible larvae indicate at least one GA-R allele at this locus was necessary for resistance in our screening bioassay at 10 μg Cry1Ac per cm^2^ diet (Table 1). However, 27% of susceptible larvae were homozygous for the GA-R allele at marker 4 (Table 1). Together these results suggest that the *r1* allele was necessary but not sufficient for resistance to Cry1Ac in our screening bioassay.

Compared to 89 control larvae reared on untreated diet, 69 resistant larvae from the F22, F23, and F26 generations had a significantly higher ratio of the GG genotype to the GL genotype for marker 4 (Fisher’s exact test; *P* < 0.0001). Based on the data for marker 4 from the F22, F23, and F26, the *r1* allele had a value for dominance *(h)* of 0.23 (Supplementary Table S3), which is intermediate between completely recessive inheritance (*h* = 0) and additive inheritance *(h =* 0.5).

To test the hypothesis that alleles at one or more other loci contributed to resistance, we compared the trajectory of resistance based on bioassay data with the trajectory of the GA-R allele at marker 4. Resistance to Cry1Ac decreased substantially over time (Figure 4; Supplementary Table S1), indicating that in the absence of Cry1Ac, a pleiotropic fitness cost was associated with one or more alleles contributing to resistance. However, the frequency of the GA-R allele at marker 4 was 0.52 in 89 control larvae from the F26, which is not different than the expected 0.50 in the F1. This suggests no fitness cost was associated with the *r1* allele, which is tightly linked with GA-R allele at marker 4. In the F26, marker 4 was in Hardy-Weinberg equilibrium as noted above, confirming the absence of selection at this locus in GA-RS. The significant decrease in resistance over time despite no decline in the frequency of the GA-R allele at marker 4 implies the decrease in resistance was caused by reduced frequency of one or more resistance alleles that carry a fitness cost in the absence of Cry1Ac and are not linked with the *r1* allele.

**Figure 4.**
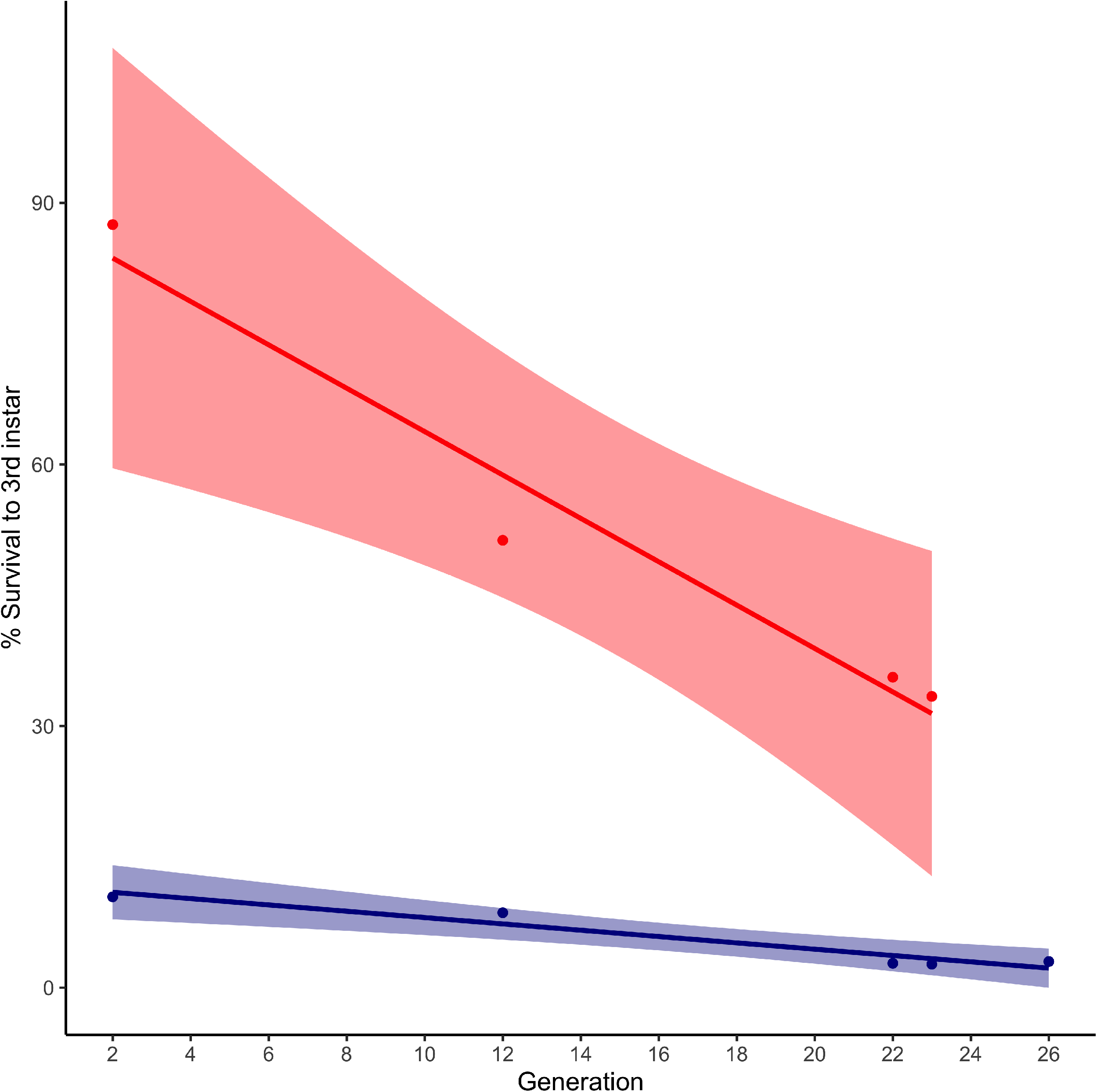
Survival of the heterogeneous GA-RS strain of *H. zea* tested on diet with 1 (red) or 10 (blue) μg Cry1Ac per cm^2^ diet. Survival at each test concentration decreased significantly. Regressions of percent survival to third instar on generation: y = −2.49x + 88.63, R^2^ = 0.94, df = 2, *P* = 0.021 and y = −0.36x + 11.67, R^2^ = 0.91, df = 3, *P* = 0.0074 for 1 and 10 μg Cry1Ac per cm^2^ diet, respectively. Generation 26 was tested only at the higher concentration. Shaded areas show 95% confidence intervals.

### Analysis of gene expression using RNA-seq

To test the hypothesis that differential gene expression contributes to resistance, we used RNA-seq to compare transcript levels between GA-R and LAB-S and between the GG and LL genotypes within GA-RS. After filtering, we analyzed expression of 12,965 genes (Supplementary Table S7). We found 2,173 differentially expressed (DE) genes between the unrelated strains LAB-S and GA-R (Supplementary Table S8) and 23 DE genes between the GG and LL genotypes within GA-RS (Supplementary Table S9). None of the genes associated with *r1* in chromosome 13 differed significantly in expression between GA-R and LAB-S or between GG and LL.

Twelve of the 23 DE genes between GG and LL were also among the set of 2,173 DE genes in the parental strain comparison, of which nine were DE in the same direction in both comparisons (higher expression in GA-R than LAB-S and in GG than LL or vice versa; Tables S6 and S7). The overlap in DE genes between these two datasets is significantly greater than expected by chance (χ^2^ = 3.86; *P* = 0.025), implying the within-strain comparison between GG and LL reflects meaningful differences between the parental strains. However, none of the 23 DE genes between GG and LL (Supplementary Table S9) are among the 11 genes previously implicated in resistance to Cry1Ac in lepidopterans (Table 3). One gene significantly downregulated in both GA-R and GG is on chromosome 1 and encodes a sodium/potassium/calcium exchanger (Supplementary Tables S8 and S9). This transmembrane protein has some functional similarities to ABC transporters and could be a candidate as a Bt receptor. However, expression was reduced only 2.6-fold in GG versus LL and 2.5-fold in GA-R versus LAB-S. Together these results indicate the *r1* region exerts a *trans*-regulatory effect on gene expression, but the current data provide no compelling evidence that any difference in expression influences resistance.

**Table 3.** GWAS and RNA-seq results for 11 genes previously implicated in lepidopteran resistance to Cry1Ac.

| Gene | Key reference | *H. zea* genome location<sup>a</sup> | Gene ID<sup>b</sup> | GWAS: *P*-value<sup>c</sup> | RNA-seq: GA-R vs. LAB-S *P*-value<sup>d</sup> | RNA-seq: GG vs. LL *P*-value<sup>d</sup> |
| --- | --- | --- | --- | --- | --- | --- |
| *ABCC1* | Chen *et al.* 2019a | 12: 9.24-9.29 | 7445 | 0.16 | **0.0014** | 0.29 |
| *ABCC2* | Baxter *et al.* 2011 | 15: 7.07-7.09 | 9800 | 0.35 | 0.95 | 0.56 |
| *APN1* | Zhang *et al.* 2009 | 9: 11.33-11.37 | 5943 | 0.53 | 0.32 | 0.33 |
| *Cadherin* | Gahan *et al.* 2001 | 6: 1.89-1.97 | 4086 | 0.46 | 1.0 | 0.13 |
| *Cadherin-86C* | Fritz *et al.* 2020 | 12: 4.60-4.62 | 7702 | 0.49 | 0.69 | 0.75 |
| *MAPK4* | Guo *et al.* 2015 | 12: 6.80-6.82 | 9824 | 0.21 | 0.12 | 0.43 |
| *mALP* | Jurat-Fuentes *et al.* 2011 | 8: 10.41-10.43 | 4895 | 0.37 | 0.11 | 0.62 |
| *Polycalin* | Wang *et al.* 2020 | 25: 2.72-2.73 | 16287 | 0.48 | 0.73 | 0.14 |
| *SP2* | Rajagopal *et al.* 2009 | 7: 1.34-1.35 | 4260 | 0.53 | 0.85 | 0.88 |
| *Tetraspanin1* | Jin *et al.* 2018 | 10: 11.58-11.59 | 6160 | 0.087 | 0.061 | 0.20 |
| *TryR* | Liu *et al.* 2014 | 27: 2.04-2.09 | 17235 | 0.60 | 0.97 | 1.0 |

<sup>a</sup> Chromosome: Mb
<sup>b</sup> Full Gene ID begins with MSTRG.
<sup>c</sup> Lowest *P*-value for any SNP in the gene based on the G' analysis
<sup>d</sup> *P*-values are FDR-corrected, bold indicates significant at < 0.05.

### Analysis of 11 genes previously implicated in lepidopteran resistance to Cry1Ac

We used our results from GWAS and RNA-seq to evaluate potential contributions to resistance of 11 genes previously implicated in lepidopteran resistance to Cry1Ac (Table 3). None of these candidate genes are in the resistance-associated QTL on chromosome 13 or the putative minor effect QTL on chromosome 10 (Table 3). In addition, none of them had any SNPs that were significantly associated with resistance in the GWAS (Table 3). Although *tetraspanin-1* is on chromosome 10 in *H. zea*, it is outside the regions of this chromosome that were moderately associated with resistance in the GWAS. As noted above, none of the 23 DE genes between GG and LL are among the 11 candidate genes (Supplementary Table S9). Only one of the 11 pairwise comparisons between strains based on RNA-seq showed a significant difference. Expression of *ABCC1* was significantly lower in GA-R than LAB-S (*P* = 0.0014, Table 3; Table S6). However, within GA-RS, expression of *ABCC1* did not differ significantly between GG and LL (*P* = 0.29; Table 3), which indicates reduced expression of *ABCC1* was not genetically linked with resistance conferred by *r1*.

## Discussion

We report three key results demonstrating a novel genetic basis of Cry1Ac resistance in the GA-R strain of *H. zea*, which resulted from field selection followed by lab selection (Brévault *et al.* 2013; Welch *et al.* 2015). First, resistance was associated with a 250-kb region of chromosome 13 that contains no genes with a previously identified role in Bt resistance or toxicity. Second, within this region, resistance to Cry1Ac was associated with a point mutation that introduces a premature stop codon in a novel candidate gene, *kinesin-12*. Third, we report evidence that one or more other loci also contributed to resistance to Cry1Ac. To facilitate these advances, we built the first chromosome-level genome assembly for *H. zea*, adding to a growing set of highly contiguous genomes for lepidopteran pests (Chen *et al.* 2019b; Ward *et al.* 2021; Yan *et al.* 2021). This genome was essential for the genetic mapping reported here and will serve as a resource for other genomic investigations into the biology of *H. zea.*

Our findings add a new candidate gene to the diverse list of genes associated with Bt resistance in lepidopterans (Jin *et al.* 2018; Guo *et al.* 2021; Jurat-Fuentes *et al.* 2021). However, the novel genetic basis of resistance does not necessarily imply a novel biochemical mechanism of resistance. The effects of *r1* could be mediated by either decreased toxin activation or reduced binding of Cry1Ac to larval midgut membranes, which are well known mechanisms of Bt resistance (Peterson *et al.* 2017; Jurat-Fuentes *et al.* 2021). Indeed, previous studies of *H. zea* have found decreased protoxin activation in GA-R (Zhang *et al.* 2019) and reduced binding of Cry1Ac to larval midgut preparations in strains with resistance caused by knocking out the putative receptor ABCC2 (Perera *et al.* 2021).

The location of *r1* on chromosome 13 is noteworthy because it corresponds closely to the region under the strongest selection in *H. zea* populations from Louisiana that were exposed to Bt crops over the past 19 years (Taylor *et al.* 2021). Although Taylor *et al.* (2021) identified a narrow region near but not containing *r1* as the most likely site of selection (~4.0 Mb in our assembly), the broader region associated with the selective sweep in their data includes *r1* (~3.8 to 5.8 Mb) and aligns with both our original and refined QTL for resistance. Thus, *r1* might be associated with resistance to Cry1Ab (which is similar to Cry1Ac) in the field populations of *H. zea* from Louisiana studied by Taylor *et al.* (2021), as well as in GA-R and its parent strain GA (Fritz *et al.* 2020), which originated from a field-selected population in Georgia in 2008 (Brévault *et al.* 2013).

The RNA-seq evidence does not support the hypothesis that altered transcription in the *r1* region causes resistance. Thus, a protein-coding mutation is more likely to be responsible for the contribution of the *r1* region to resistance. We hypothesize that this contribution is mediated by the premature stop codon in the *kinesin-12* gene. Among the protein-coding mutations in the candidate region, only this nonsense mutation that shortens the predicted protein by 40% is expected to have a major effect on protein function. Furthermore, of the 10 genes in the region tightly associated with resistance, midgut expression was highest for *kinesin-12*, suggesting a midgut function for the protein it encodes. Protein functional prediction algorithms including DeepGoPlus provide moderate support for the original annotation as a kinesin with a function in intracellular transport or structure. Nonetheless, we do not know the normal function of the kinesin-12 protein and cannot infer that the C546T mutation causes resistance. In future work, we aim to test the hypothesis that the C546T mutation contributes to resistance by determining if resistance is reduced when we use CRISPR/Cas9 to replace the mutant sequence in GA-R with the wild type sequence from LAB-S (Jin *et al.* 2018; Fabrick *et al.* 2021). If disruption of the *kinesin-12* gene is not sufficient for resistance, as the results here imply, we expect that knocking out this gene would not cause resistance in a susceptible strain.

Kinesins and kinesin-related proteins are motor proteins important in microtubule function, chromosomal movement, and organelle transport (Ali and Wang 2020) that have not been associated previously with Bt resistance. Several kinesins are involved in mitogen-activated protein kinase (MAPK) signaling cascades (Liang and Yang 2019) and MAPK signaling is implicated in Bt resistance (Guo *et al.* 2015, 2020, 2021; Qin *et al.* 2021). Furthermore, a case of xenobiotic resistance in mice involved a mutant kinesin acting downstream of a MAPK (Watters *et al.* 2001). Thus, one hypothesis is that *kinesin-12* acts downstream of MAPK as part of the signaling cascade initiated when Cry1Ac binds to a midgut receptor. However, MAPK influences Bt resistance via downregulation of Bt receptors (Guo *et al.* 2015, 2020, 2021; Qin *et al.* 2021). Here we see no evidence for reduced transcription of putative receptors, making this an unlikely explanation for the link between *kinesin-12* and resistance.

Kinesins also play a role in the localization of transmembrane proteins to the cell membrane (Jana *et al.* 2021) and in intracellular cadherin trafficking (Phang *et al.* 2014). The transport functions of kinesins and kinesin-related proteins entail motor complexes of three or more proteins (Phang *et al.* 2014), suggesting interactions between different proteins could be interrupted to interfere with receptor transport to the membrane. We hypothesize that in the GA-R strain of *H. zea*, a truncated *kinesin-12* in combination with mutations affecting one or more of its interacting partners blocks proper localization of a Bt receptor on the membrane.

The results from GWAS and fine-scale mapping show the *r1* allele was necessary, but not sufficient for resistance in our screening bioassay, implying contributions from one or more additional loci. If a second unlinked mutation were also necessary for resistance in our screening bioassay, this would be expected to yield a second major peak in the GWAS, similar to the peak for the QTL in chromosome 13. The lack of a second major peak suggests that mutations in two or more unlinked loci could each cause resistance in combination with *r1* (e.g., *r1* plus mutation *X* or *r1* plus mutation *Y* could cause resistance).

The decline in resistance over time in GA-RS also implies more than one locus contributed to resistance. While resistance to Cry1Ac declined significantly across generations in GA-RS, the frequency of C546T and other *r1* linked alleles did not. After 22, 23 and 26 generations without exposure to Bt toxins, it was not lower than its expected initial frequency of 0.50. Thus, the decline in resistance reflects a decreased resistance allele frequency at one or more other loci. Unlike the C546T mutation, which did not have a substantial fitness cost in the lab, the observed decline in resistance suggests that a fitness cost was associated with at least one mutation at another locus that contributed to resistance in GA-RS. Polygenic resistance to Cry1Ac or Cry1Ab also has been reported in strains of *H. zea* unrelated to GA-R (Caccia *et al.* 2012; Lawrie *et al.* 2020; Perera *et al.* 2021; Taylor *et al.* 2021) and in other species of Lepidoptera (Kaur and Dilwari 2010; Zhao *et al.* 2021; Ma *et al.* 2022).

The results summarized above have implications for understanding the trajectory of the C546T mutation in the field. The high frequency of the C546T mutation in the field-selected GA strain that was started with 180 field-collected larvae suggests this mutation was common in 2008 in the moderately resistant field population in Georgia from which GA was derived (Brévault *et al.* 2013). In the absence of exposure to Cry1Ac, the frequency of this mutation did not increase significantly in GA from the F72 to F87 or in GA-RS from the expected frequency in the F1 to the observed frequency in the F26. Thus, because GA was not exposed to Cry1Ac in the lab, it is unlikely the frequency of this mutation was low initially in GA and subsequently increased because of strong selection. Nonetheless, we did not detect this mutation in Cry1Ac-resistant populations from the same site in Georgia in 2021 or in Arizona in 2020. Thus, this mutation is not associated with resistance to Cry1Ac in all field populations of *H. zea.* Also, its frequency apparently decreased in the field in Georgia from 2008 to 2021. One hypothesis is that the frequency of this mutation decreased in Georgia because it has a substantial fitness cost under field conditions, such as reduced overwintering survival (Carrière *et al.* 2001), which would not be evident in the lab. The C546T mutation could have been replaced by one or more mutations that have lower fitness costs (Guillemaud *et al.* 1998), confer higher resistance to Cry1Ac, and/or confer resistance without contributions from mutations at other loci. More research is needed to determine the function of kinesin-12 and its role in resistance, as well as the genetic basis of Cry1Ac resistance in current field populations of *H. zea.*

## Data and code availability

All raw sequence data is available at NCBI (Bioproject: PRJNA767434). Phenotypic data, HRM and Sanger genotyping data, initial and final genome assemblies, genome annotations, and scripts for analyses and figures are all available via OSF. Supplementary materials are available at figshare.

## Acknowledgments

Mention of trade names or commercial products in this article is solely for the purpose of providing specific information and does not imply recommendation or endorsement by the U.S. Department of Agriculture. USDA is an equal opportunity provider and employer. We thank Xinzhi Ni for sending *H. zea* from Georgia; Alex Yelich and Chandran Unnithan for help with insect rearing and dissections; Yidong Wu, David Heckel, Megan Fritz, Katherine Taylor, Fred Gould, and Juan Luis Jurat-Fuentes for their valuable comments on the manuscript; and Jon Galina-Mehlman, Jayson Talag, and Dave Kudrna for their assistance and advice regarding PacBio and Illumina sequencing.

## Author contributions

J.F., Y.C., B.E.T., and L.M.M. conceived and designed the project. K.M.B., C.W.A., B.A.D., and X.L. performed experiments and collected the data. K.M.B., C.W.A., Y.C., and B.E.T. analyzed data. K.M.B. and B.E.T. wrote the manuscript with input from all authors.

