## Supplementary Figures and Tables for "Novel genetic basis of resistance to Bt toxin Cry1Ac in *Helicoverpa zea*"

**Supplementary Table S1** Resistance to Cry1Ac in the GA-RS strain of *Helicoverpa zea* based on 7-day diet bioassays.

| GA-RS generation | Concentration (µg Cry1Ac per $cm^2$ diet) | Number of neonates tested | Percentage (%) | | | |
|---|---|---|---|---|---|---|
| | | | Dead | First instar | Second instar | ≥ Third instar |
| 2 | 1 | 48 | 4.2 | 0.0 | 8.3 | 87.5 |
| 2 | 10 | 48 | 18.8 | 22.9 | 47.9 | 10.4 |
| 12 | 1 | 3839 | 6.1 | 9.4 | 33.2 | 51.3 |
| 12 | 10 | 3712 | 34.9 | 22.8 | 33.7 | 8.6 |
| 22 | 1 | 640 | 11.2 | 16.2 | 36.9 | 35.6 |
| 22 | 10 | 1280 | 62.8 | 14.8 | 19.6 | 2.8 |
| 23 | 1 | 1274 | 11.1 | 14.4 | 41.1 | 33.4 |
| 23 | 10 | 1274 | 58.7 | 20.0 | 18.6 | 2.7 |
| 26 | 10 | 3788 | 63.2 | 17.9 | 16.0 | 3.0 |

We analyzed generation 12 using GWAS and generations 22, 23, and 26 with fine-scale mapping. For control larvae reared on diet without Cry1Ac, mean survival to third instar was 96% (range: 91-100%, mean n = 99 control larvae per bioassay in five bioassays).

**Supplementary Table S2** Primers used to amplify HRM genotyping sites. The K12 site was used to genotype at the C564T mutation causing a stop codon in *kinesin-12*.

| Site | Forward Primer (5` - 3`) | Reverse Primer (5` - 3`) |
|---|---|---|
| 1 | CACAAATGTTCTGTGAGTCAAT | TAATCAAGGAAGTGTGCAAGA |
| 2 | GGTAGAAAGCTGGCAATCA | GGAAGAAGAACTCGACAGTG |
| 3 | GCCAACCAGTTCCATCTC | GCAACTTCAATCGTACTAATGTC |
| 4 | TTTAAACCGACAGTAGTACAGG | CATCCGACACGCAACAA |
| 5 | GTCACCGTCCATATCTTCTTC | TGGAGAACCTTCAGTAAACAC |
| 6 | CGTTCTTATATTTCATCCGCAA | TATTACGTAAGAGACCGCAAA |
| 7 | GGTTTAGTCATTACCCGTGTT | CTCATTAGGAGCGGTAGTTTAG |
| 8 | GGAAGCTTGCCAGACAAA | CTATTTACGTAGAGAAGGCTGTC |
| 9 | GCCACAATTAAATTAACGACCA | TGAATGACTATTACGACACAGC |
| 10 | CTAGACACCTTGCGAATGAC | CGCCTACAATAGGACATTACAT |
| 11 | CTGTCGAAGTTACTGTAGTTATCAT | TCTTTGTGTCATTACATTGGTG |
| 12 | TGGATGACGTAATCGTGTG | CCGTTTCAACTGAACACTTC |
| K12 | CCGAACTTGAGGCAAGA | GTTACCCGTGCTGTCTTT |

**Supplementary Table S3** Calculation of dominance (*h*) of resistance based on genotypes at marker 4 for *H. zea* larvae from GA-RS in the F22, F23 and F26 generations.

| | Genotype at marker 4 | | | | |
|---|---|---|---|---|---|
| Generation | GG | GL | LL | GL/GG | *h* |
| *Resistant larvae* | | | | | |
| F22 & F23 | 40 | 17 | 0 | 0.425 | |
| F26 | 16 | 7 | 0 | 0.438 | |
| Total | 56 | 24 | 0 | 0.429 | |
| *Control larvae* | | | | | |
| F26 | 24 | 44 | 21 | 1.833 | 0.23 |
| *Susceptible larvae* | | | | | |
| F22 & F23 | 13 | 24 | 12 | 1.846 | 0.23 |

G indicates the allele was more common in the resistant GA-R strain, L indicates the allele was more common in the susceptible LAB-S strain. At marker 4, which is relevant here, allele G was at 100% in GA-R and L was at 100% in LAB-S, so we can infer G originated from GA-R and L from LAB-S (see text).

We calculated the dominance parameter *h*, which varies from 0 for recessive resistance to 1 for dominant resistance, based on the equation from Liu and Tabashnik (1997):

(1) $h = (w_{12} - w_{22})/(w_{11} - w_{22})$

where $w_{11}$, $w_{12}$, and $w_{22}$ are the fitness values at a particular toxin concentration for resistant homozygotes, heterozygotes, and susceptible homozygotes, respectively. Typically, survival at the given toxin concentration is used to estimate fitness, which is reasonable because the survival of each genotype is likely to be correlated with its fitness.

To calculate *h*, we used the data from marker 4 to estimate survival of putative resistant homozygotes (GG), heterozygotes (GL), and susceptible homozygotes (LL) exposed to 10 micrograms Cry1Ac per $cm^2$ diet. At this concentration, survival of LL was 0.

When survival of susceptible homozygotes = 0, equation (1) reduces to:

(2) $h = w_{12}/w_{11}$.

Thus, we estimated *h* as the survival of GL relative to GG for larvae exposed to diet with 10 micrograms Cry1Ac per $cm^2$ diet. We calculated *h* as the ratio of GL/GG for resistant larvae (which survived on diet on with 1 microgram Cry1Ac per $cm^2$ diet) divided by the ratio of GL/GG for control or susceptible larvae. For resistant larvae, the number of GL relative to GG did not differ significantly between F22 and F23 versus F26 (Fisher's exact test, $P = 1$). Thus, we used the ratio of GL/GG from pooling data from F22, F23, and F26 (0.429) to calculate *h*.

In principle, GL/GG in GA-RS is best estimated by GL/GG for control larvae, which were reared on untreated diet. However, GL/GG was similar for control larvae (1.833) and susceptible larvae (1.846), which were live first instars on diet with 1 microgram Cry1Ac per $cm^2$ diet. Therefore, the value of *h* was 0.23 using GL/GG in GA-RS calculated from either control or susceptible larvae. We note that the dominance of *r1* could be affected by variation in the frequency of one or more resistance alleles at other loci.

**Supplementary Table S4** Genome assembly and annotation statistics for the *de novo H. zea* genome as compared to that of Pearce *et al.* (2017).

| | Current assembly | Pearce et al. 2017 assembly |
|---|---|---|
| Genome size (Mb) | 375.2 | 341.1 |
| # of scaffolds | 31 | 2975 |
| Scaffold N50 length (Mb) | 12.9 | 0.2 |
| # of contigs | 32 | 34,676 |
| Contig N50 length (Mb) | 12.9 | 0.01 |
| Gaps (Mb) | 0.0001 | 34.1 |
| GC content (%) | 36.9 | 36.2 |
| Repeat content (%) | 33.0 | 16.0 |
| Number of proteins | 15,842 | 15,200 |
| Genome BUSCO (%) | | |
| Complete | 98.9 | 96.6 |
| Single copy | 98.5 | 95.4 |
| Duplicated | 0.4 | 1.2 |
| Fragmented | 0.3 | 1.1 |
| Missing | 0.8 | 2.3 |
| Proteome BUSCO (%) | | |
| Complete | 93.6 | 91.9 |
| Single Copy | 93.0 | 91.0 |
| Duplicated | 0.6 | 0.9 |
| Fragmented | 1.2 | 1.9 |
| Missing | 5.2 | 6.2 |

**Supplementary Table S5** Amino acid sequence identity for the predicted kinesin-12 protein in five species of Lepidoptera relative to the LAB-S strain of *H. zea*

| | Overall identity (%) | Identity upstream of C546T (%) | Identity downstream of C546T (%) |
|---|---|---|---|
| *H. armigera* | 97 | 98 | 97 |
| *C. virescens* | 87 | 91 | 85 |
| *S. litura* | 61 | 74 | 55 |
| *M. sexta* | 44 | 44 | 44 |
| *B. mori* | 38 | 38 | 44 |
| Mean | 65 | 69 | 65 |

**Supplementary Table S6** GO categories identified by DeepGOPlus to be associated with the kinesin-12 protein and their confidence scores. Only predictions scores above a threshold of 0.3 are shown

| GO Type | GO term | GO description | Prediction Score |
|---|---|---|---|
| Cellular Component | | | |
| | GO:0110165 | cellular anatomical entity | 0.453 |
| | GO:0043226 | organelle | 0.412 |
| | GO:0005622 | intracellular anatomical structure | 0.403 |
| | GO:0043229 | intracellular organelle | 0.381 |
| | GO:0043232 | intracellular non-membrane-bounded organelle | 0.381 |
| | GO:0043228 | non-membrane-bounded organelle | 0.381 |
| Molecular Function | | | |
| | GO:0005488 | binding | 0.358 |
| Biological Process | | | |
| | GO:0009987 | cellular process | 0.370 |

**Supplementary Table S7** Results of differential expression for all 12,965 genes analyzed in edgeR.

**Supplementary Table S8** The genes differentially expressed between GA-R and LAB-S 3rd instar midguts.

**Supplementary Table S9** The genes differentially expressed between RR and SS genotypes.

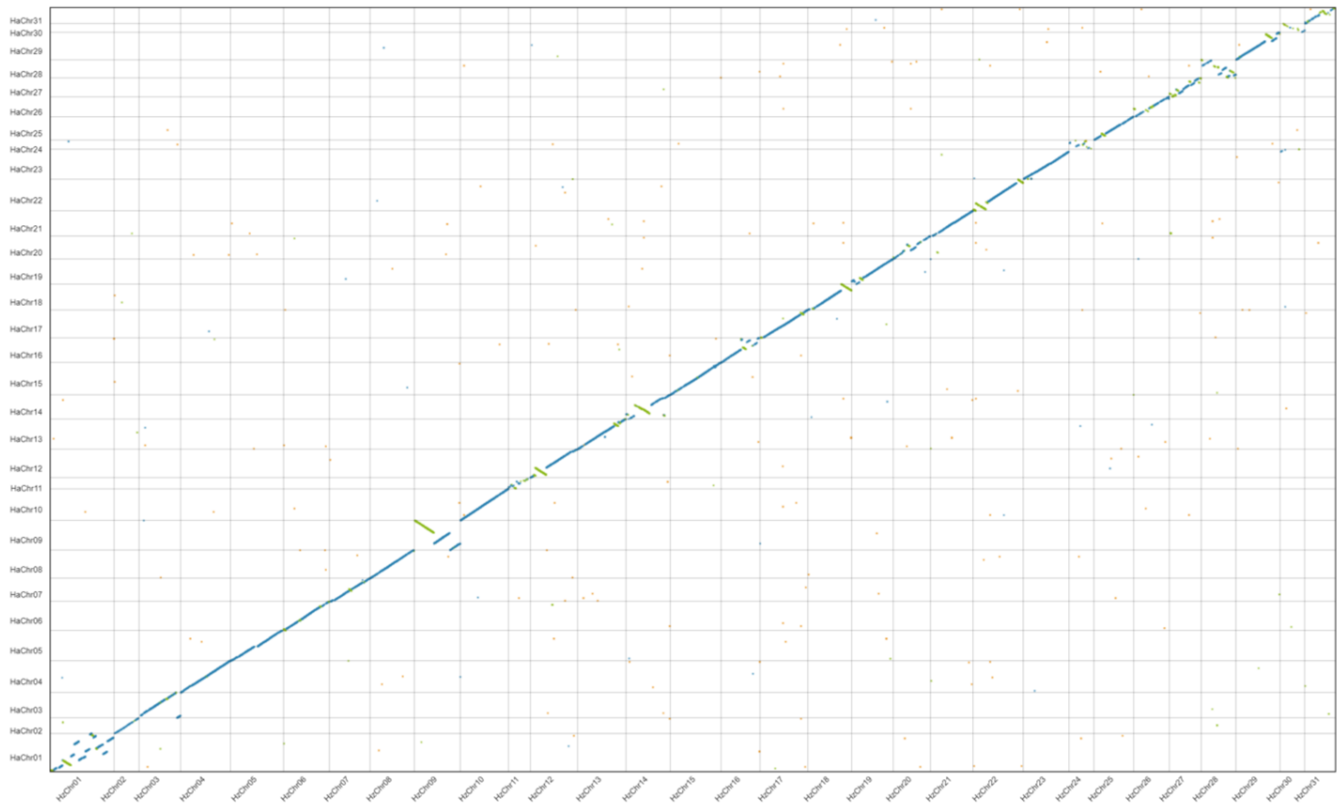

**Supplementary Figure S1** Syntenic comparisons of chromosomes from our *de novo H. zea* assembly (x-axis) and *H. armigera* (y-axis; Valencia-Montoya *et al.* 2020). Green lines that are perpendicular to the main line in blue (from the lower left to the upper right) represent chromosomal inversions. Chromosome 1, the Z chromosome, has one inversion (see text for details).

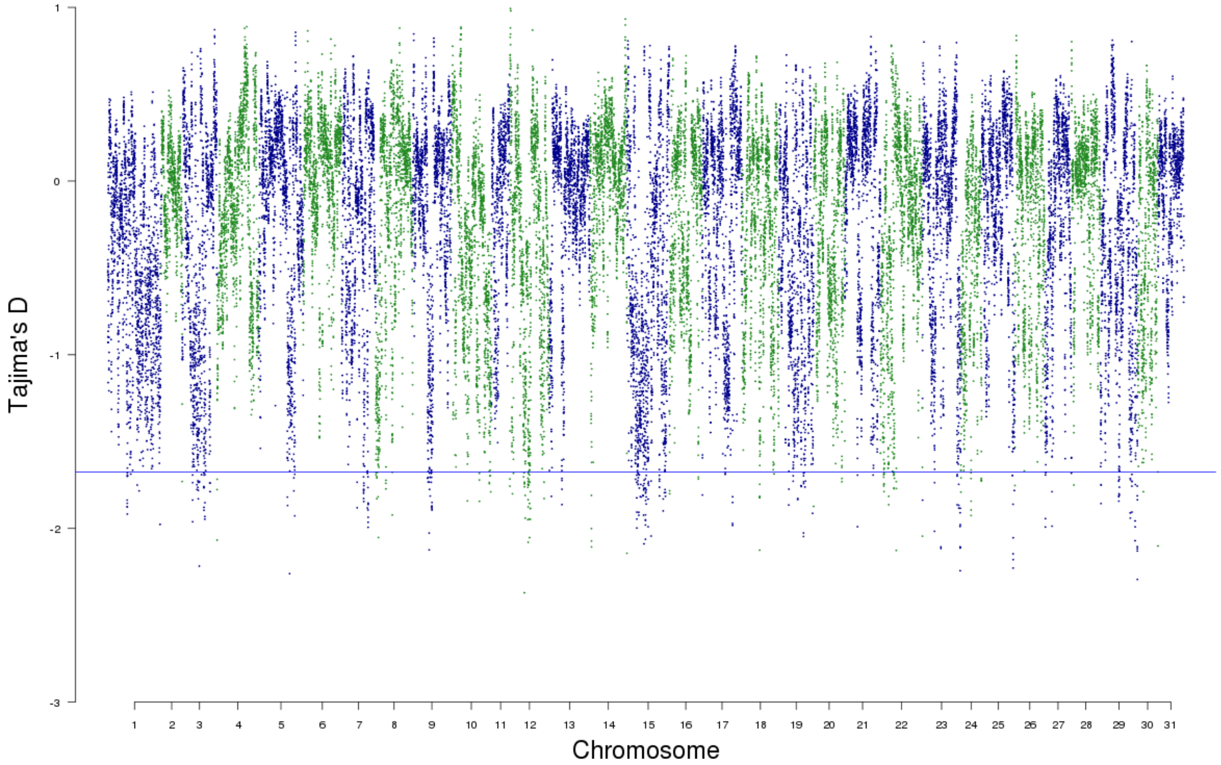

**Supplementary Figure S2** Genome wide measures of Tajima’s D across 50-kb sliding windows in GA-R. The blue line represents the 95% percentile of the distribution.

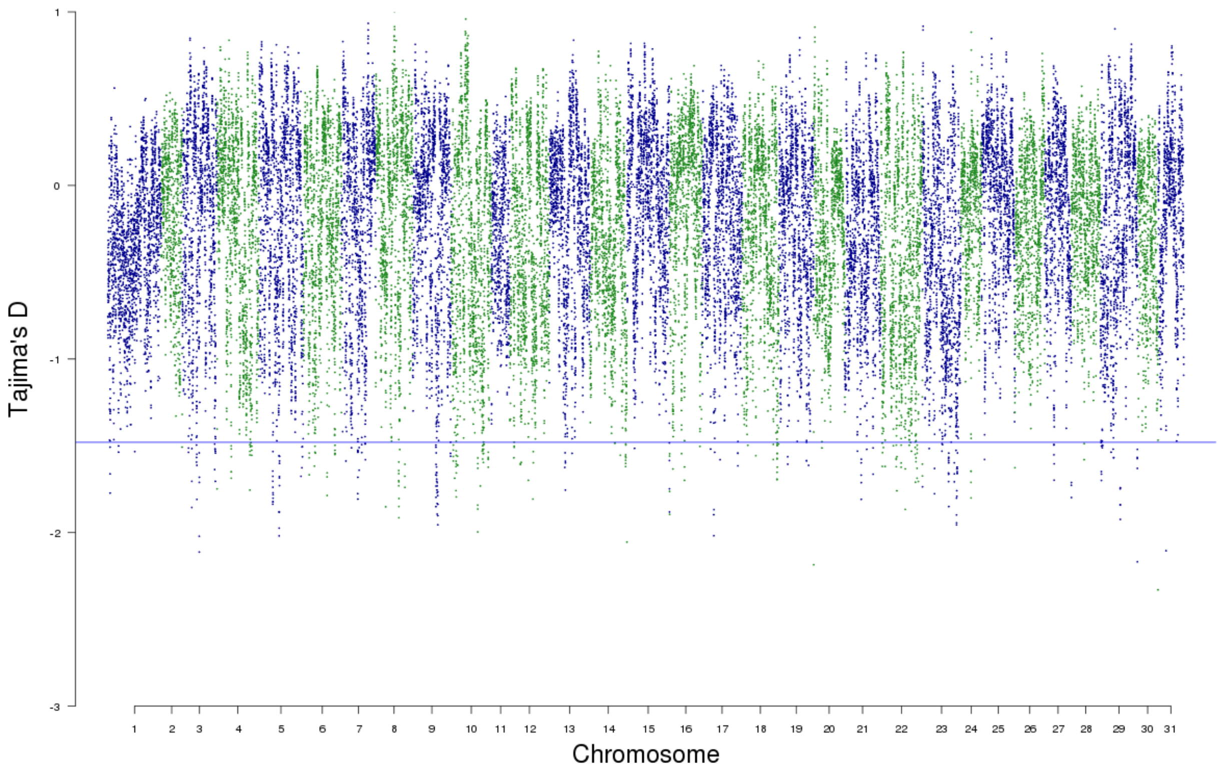

**Supplementary Figure S3** Genome wide measures of Tajima’s D across 50-kb sliding windows in LAB-S. The blue line represents the 95% percentile of the distribution.

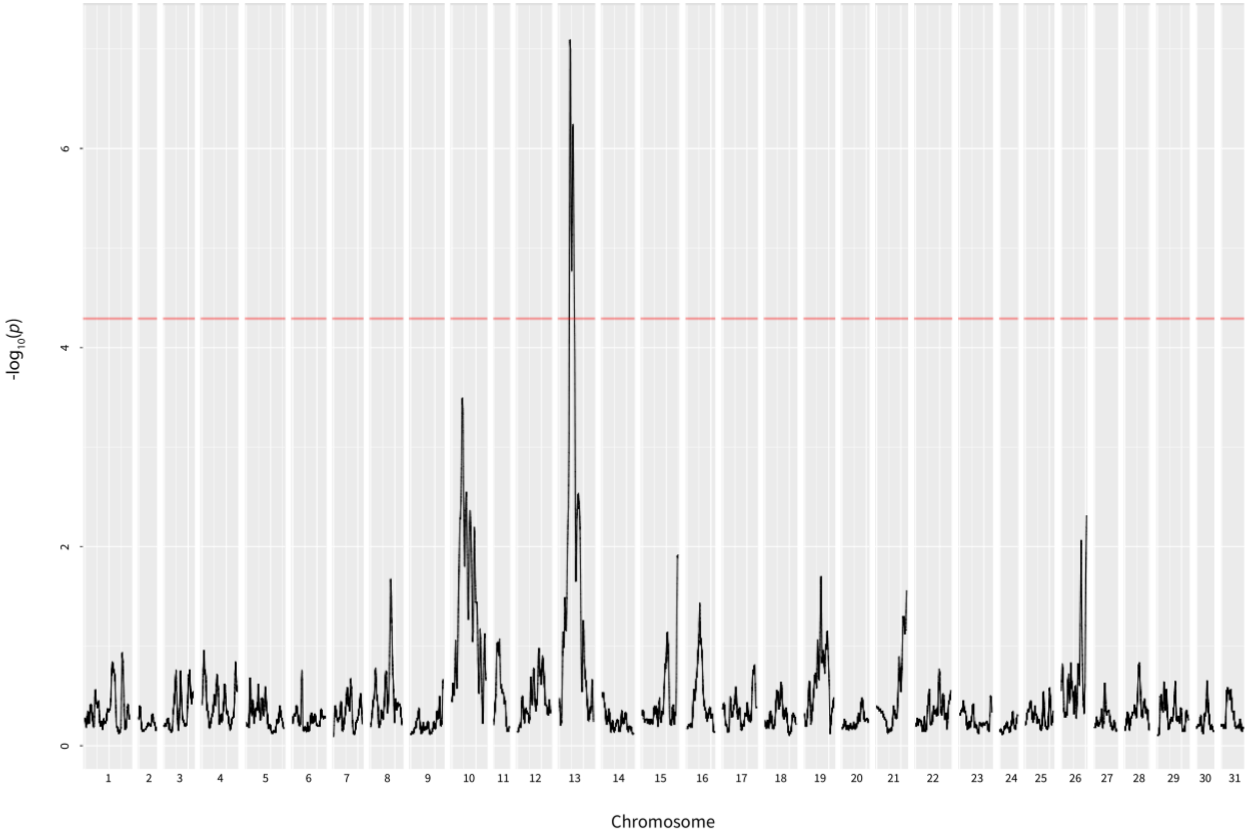

**Supplementary Figure S4** Probability values for 500-kb sliding windows from the G` analysis of the GWAS data. Only the QTL on chromosome 13 is statistically significant. The QTL on chromosome 10, which might reflect a minor effect on resistance, is not statistically significant. The red line corresponds to an FDR-corrected *P*-value of 0.05.

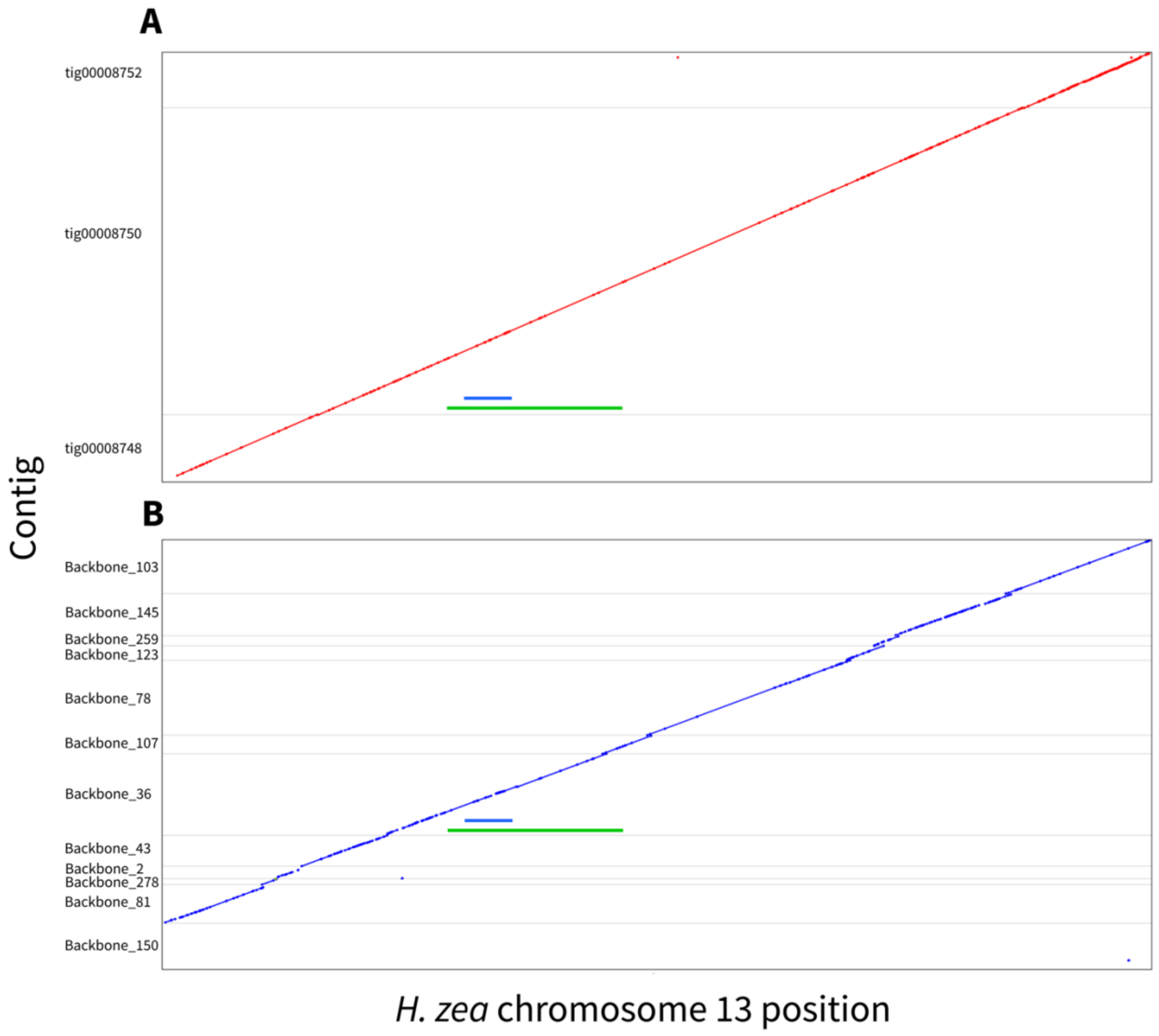

**Supplementary Figure S5** Assembled chromosome 13 contigs plotted against the broad QTL (4.0 – 6.5 Mb) in green and the region within this QTL that was tightly linked with resistance (4.3 – 4.6 Mb) in blue. (A) Contigs from the Canu PacBio assembly. (B) Contigs from the hybrid DBG2OLC assembly. The narrower region was captured in a single contig in both assemblies. The wider region was captured in a single contig in our Canu assembly. In our DBG2OLC assembly, all of the wider region except from 6.3 to 6.5 Mb, which was not closely linked with resistance, was captured in a single contig.

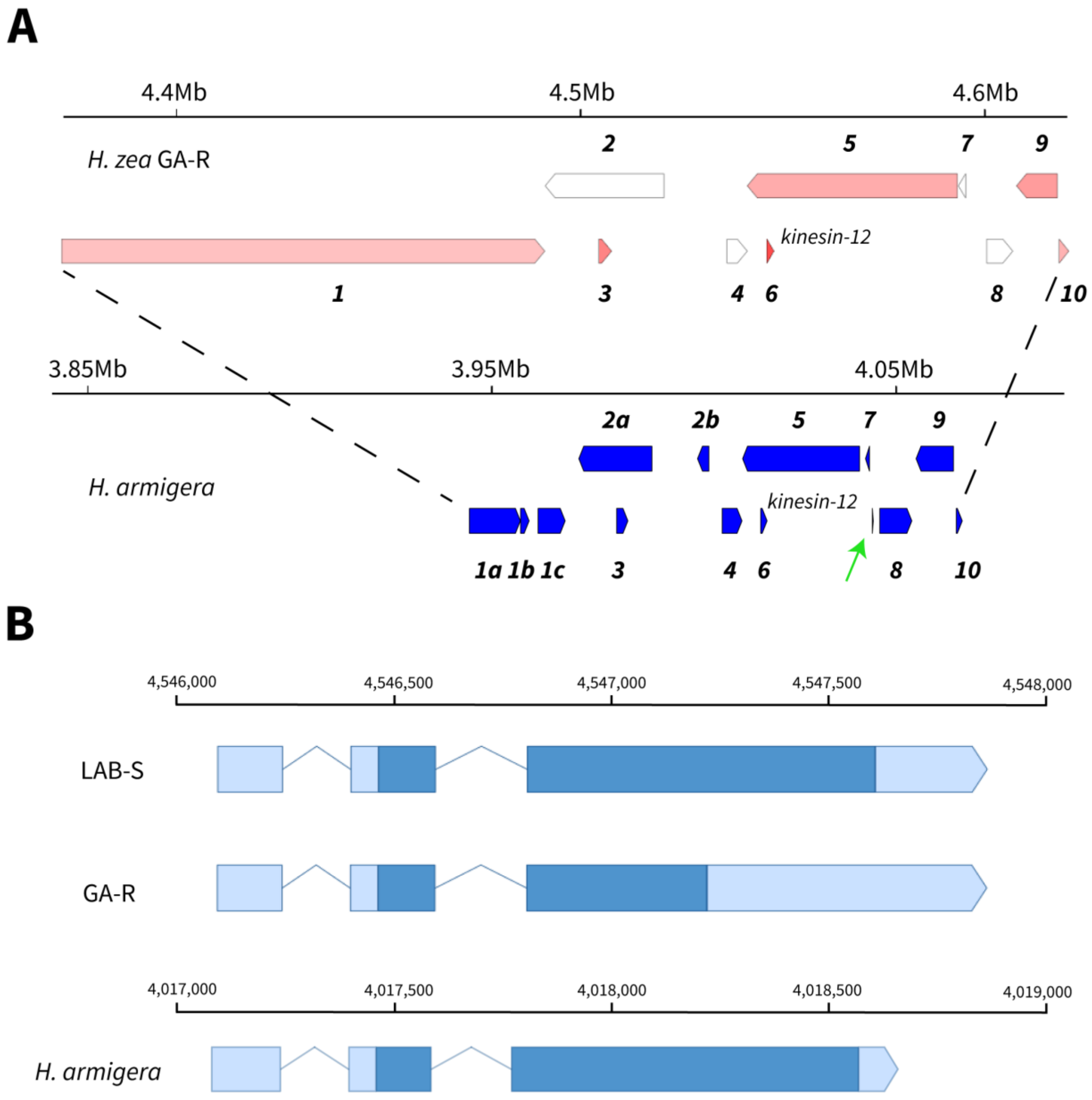

**Supplementary Figure S6** Comparison of genes between *H. zea* and *H. armigera* in the *r1* region. (A) Genes in the *r1* region. Reverse strand at top, forward strand below for each species. Genes that are homologous between species share the same number (from 1 to 10). For *H. zea*,10 genes occur in the region and seven of these (shown in pink) are expressed in 3rd instar midguts (Table 2). For *H. armigera*, 14 genes are annotated (Pearce *et al*. 2017; Valencia-Montoya *et al.* 2020); midgut expression data are available but not shown here. Of the four additional annotated genes in *H. armigera*, three have homology to parts of annotated *H. zea* genes, which suggests these differences between species reflect splitting genes in *H. armigera* versus merging them in *H. zea*. One gene in *H. armigera* (marked by a green arrow) was not annotated in *H. zea* but is only 233 bp long, and therefore unlikely to be a protein-coding gene. (B) Structure of the *kinesin-12* gene in the LAB-S and GA-R strains of *H. zea* compared with *H. armigera*

(XM_021337566.1). Dark blue represents coding regions, light blue represents UTR. The numbers above the lines represent base pair positions on chromosome 13.

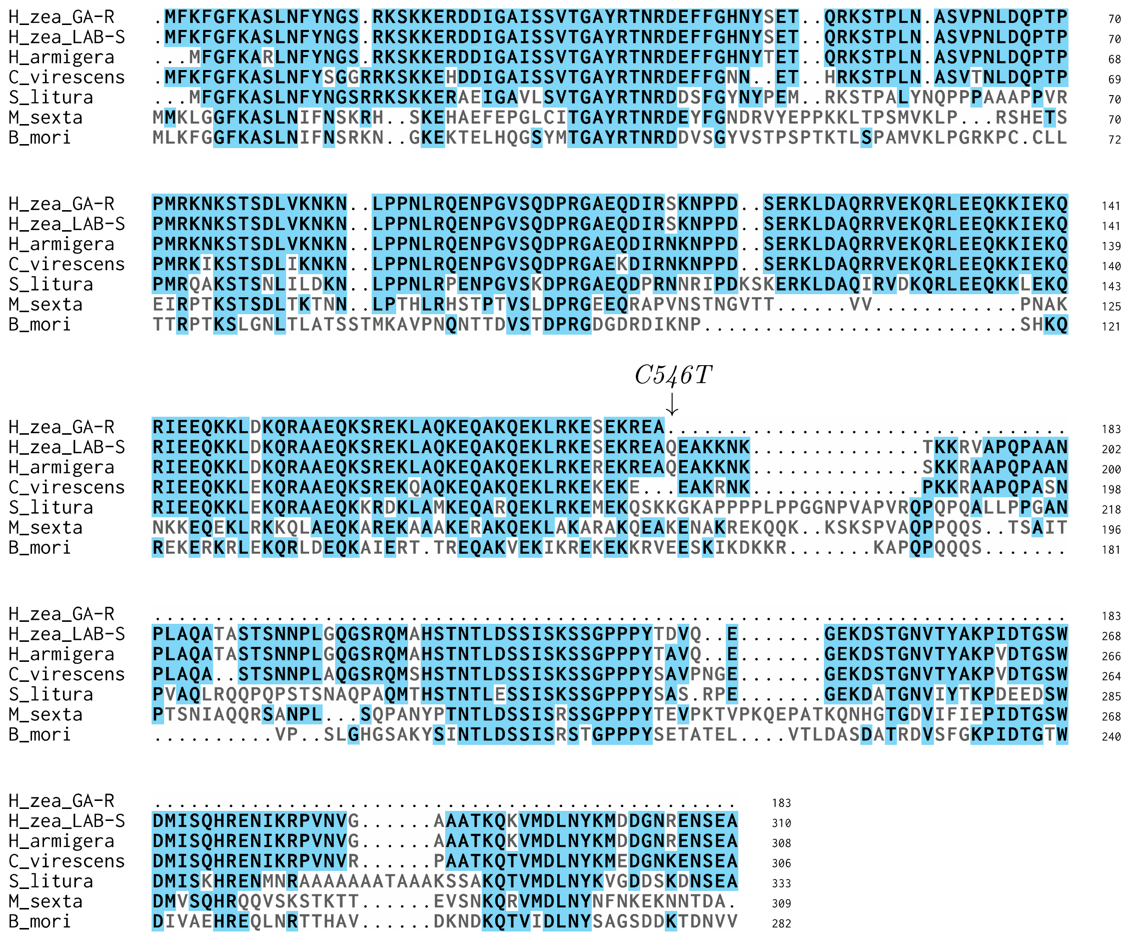

**Supplementary Figure S7** Alignments of kinesin-12 including both forms from *H. zea*, *H. armigera* (XP_021193241.1), *Chloridea virescens* (PCG76683.1), *Spodoptera litura* (XP_022828947.1), *Manduca sexta* (KAG644083.1), and *Bombyx mori* (XP_004927959). Consensus amino acids are highlighted. The location of the stop-codon mutation is marked as C546T. The annotated sequence in *B. mori* contained extra amino acids at the start of the protein, which we removed for this alignment to match the other six sequences.

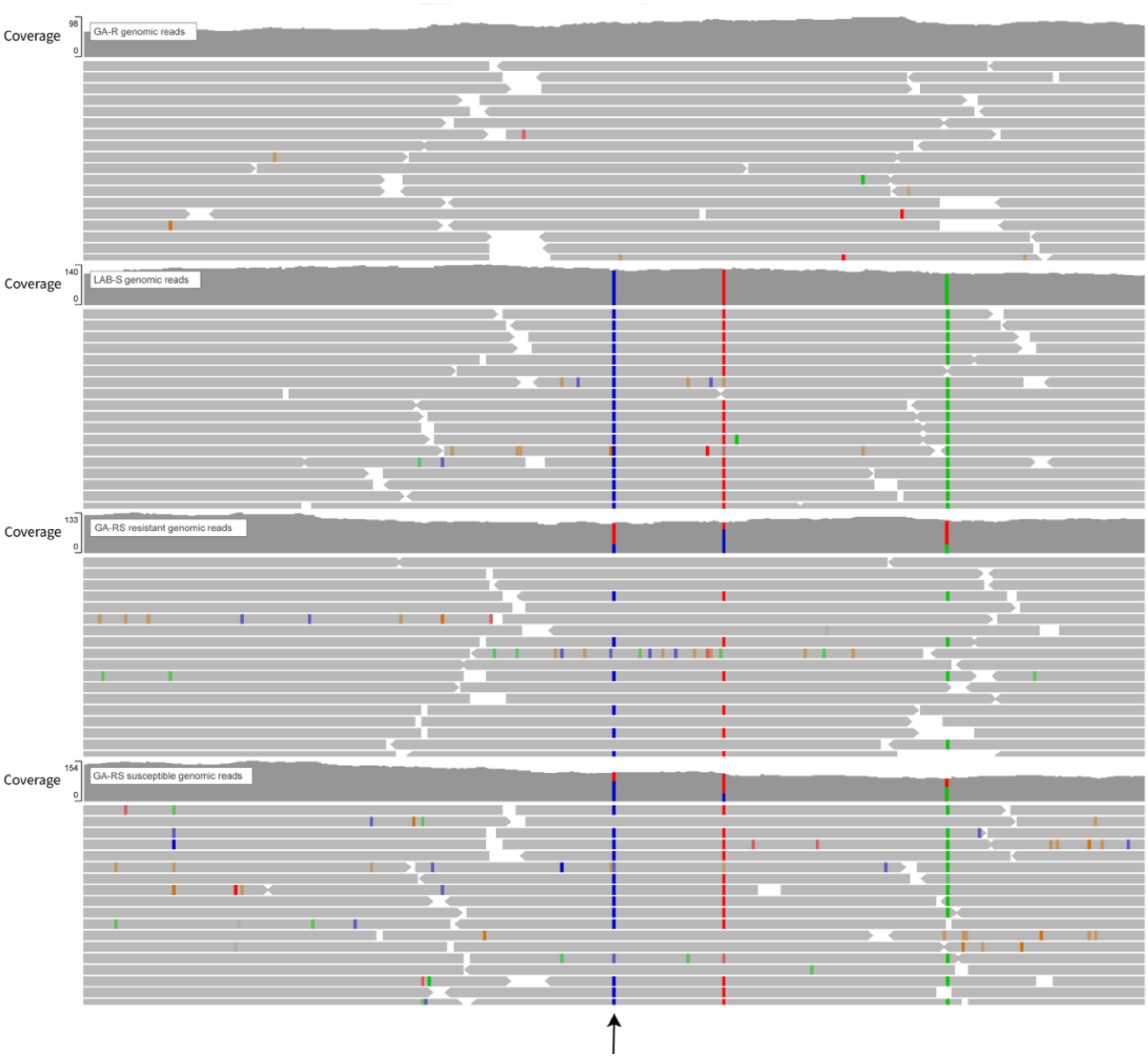

**Supplementary Figure S8** Mapped genomic reads from GA-R, LAB-S, resistant and susceptible GA-RS samples, with the arrow pointing to the *kinesin-12* stop codon mutation. Colored bars indicate reads containing SNPs different from the GA-R reference.

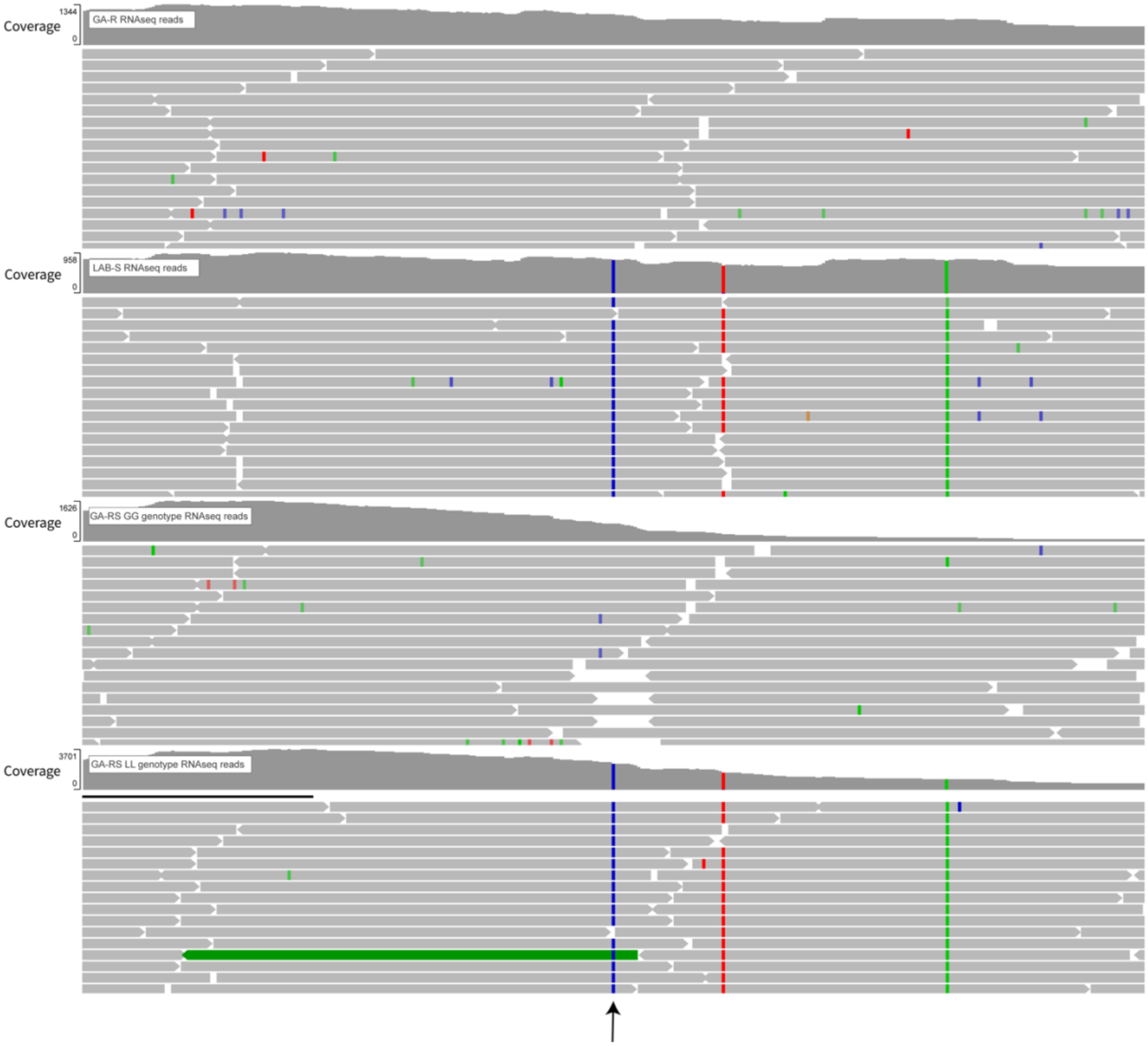

**Supplementary Figure S9** Mapped RNA-seq reads from GA-R, LAB-S, GG, and GL samples, with the arrow pointing to the *kinesin-12* stop codon causing mutation.

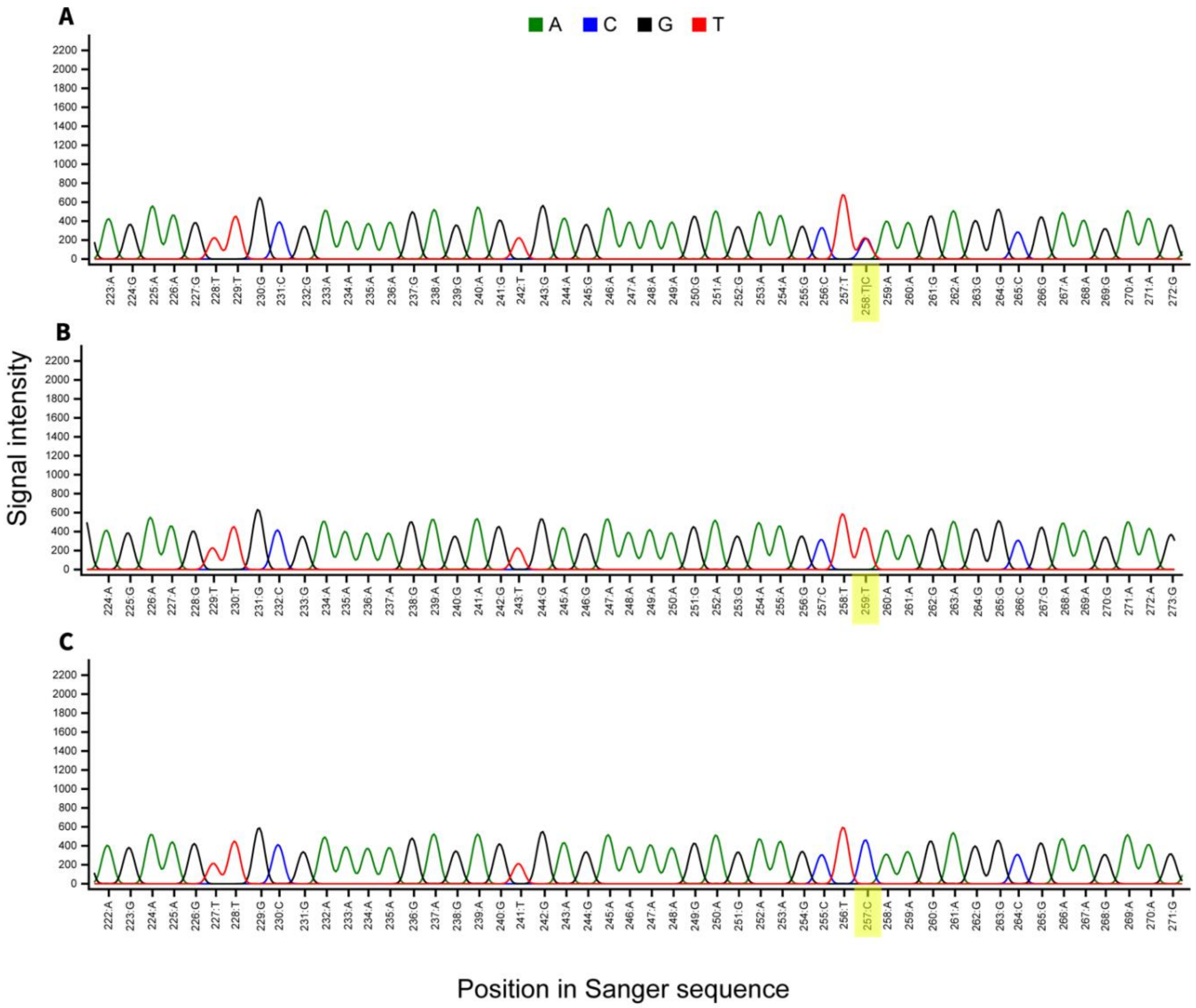

**Supplementary Figure S10** Sanger sequencing chromatograms for showing the position of the *kinesin-12* stop codon mutation highlighted in yellow. (A) A GA heterozygote sample. (B) A GA sample homozygous for the stop codon. (C) A Tifton, GA field sample homozygous for the wild type allele with no stop codon.

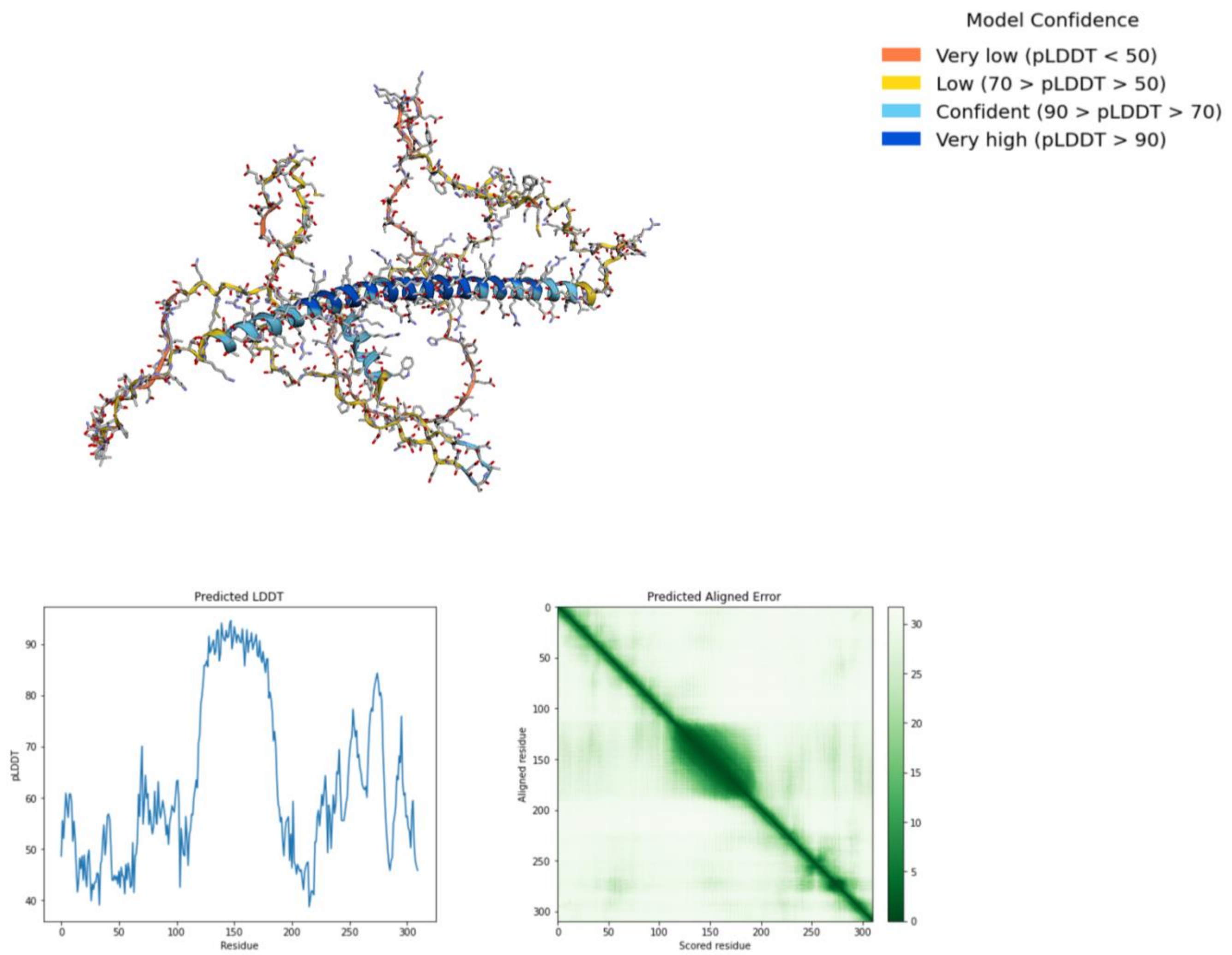

**Supplementary Figure S11** Predicted protein structure of the full-length (LAB-S) *H. zea* kinesin-12 from AlphaFold.

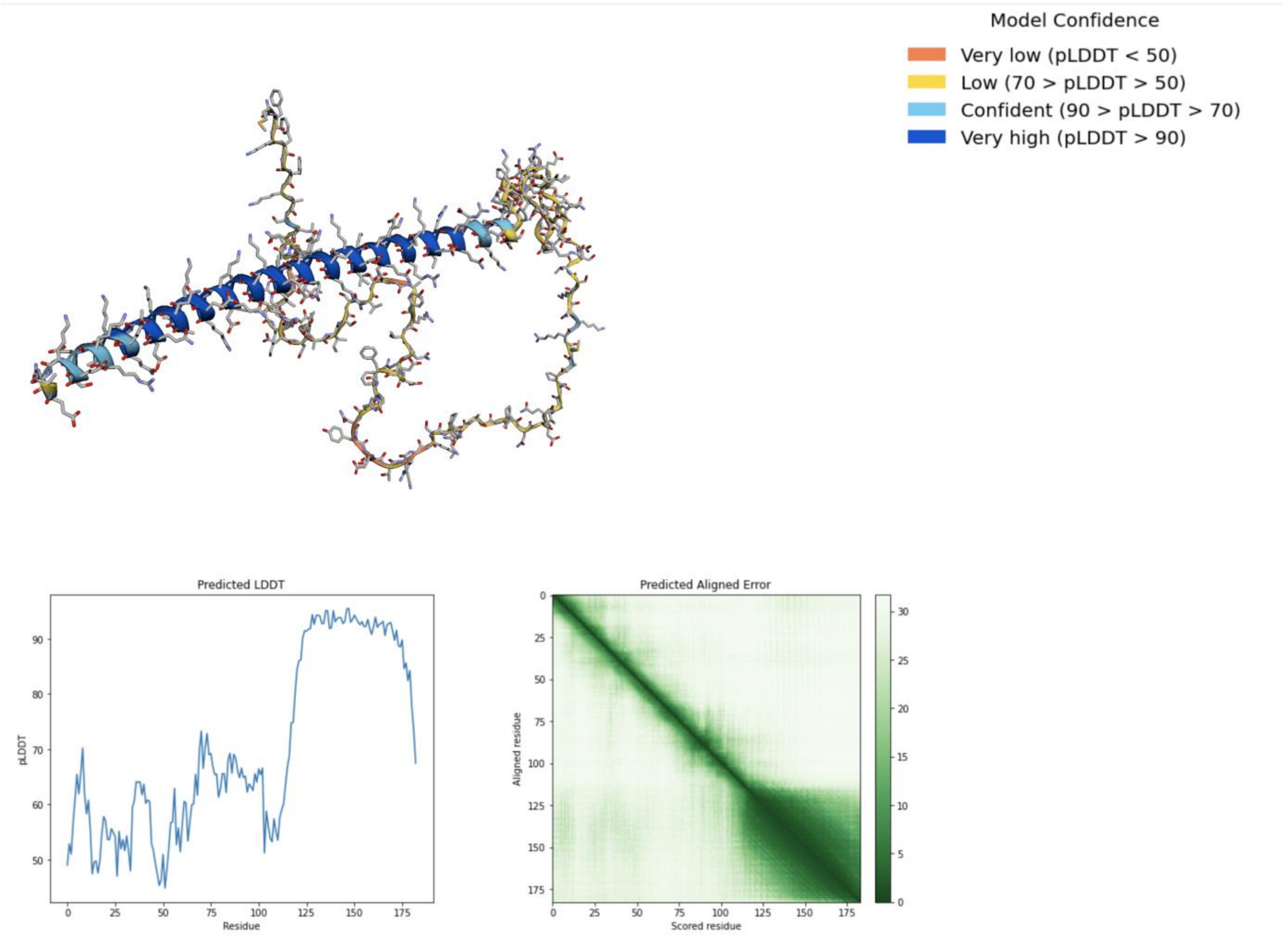

**Supplementary Figure S12** Predicted protein structure of the truncated (GA-R) *H. zea* kinesin-12 from AlphaFold.
